# Fate specification triggers a positive feedback loop of TEAD–YAP and NANOG to promote epiblast formation in preimplantation embryos

**DOI:** 10.1101/2024.08.15.608179

**Authors:** Naoki Hirono, Masakazu Hashimoto, Hinako Maeda, Hiromi Shimojo, Hiroshi Sasaki

## Abstract

In preimplantation embryos, epiblast (EPI) fate specification from the inner cell mass is controlled by the segregation of NANOG and GATA6 expression. TEAD–YAP interaction is activated during EPI formation, and is required for pluripotency factor expression. These events occur asynchronously with similar timing during EPI formation, and their relationship remains elusive. Here, we examined the relationship between NANOG–GATA6 and TEAD–YAP. The nuclear accumulation of YAP takes place only in EPI-specified cells, and a positive feedback loop operates between NANOG and TEAD–YAP. The effects of TEAD–YAP on SOX2 upregulation in EPI-specified cells are likely indirect. EPI fate specification also alters the response of *Nanog, Sox2* and *Cdx2* to TEAD–YAP. These results suggest that EPI-fate specification alters the transcriptional network from the morula-like to the EPI-specified state and activates TEAD–YAP to trigger a positive feedback loop with NANOG, which stabilizes the EPI fate. The coordinated occurrence of these processes in individual cells likely supports proper EPI formation under the condition of asynchronous EPI-fate specification.

**Summary statement:** Epiblast fate specification by segregation of NANOG from GATA6 alters the transcriptional network and activates TEAD–YAP, triggering a positive feedback loop with NANOG to promote epiblast formation.

## Introduction

Recent advances in single-cell research have revealed that cell behavior during development, e.g., division pattern, lineage, and gene expression, fluctuates and varies (Briggs et al., 2018; Farrell et al., 2018; Kurotaki et al., 2007; Morris et al., 2010; Niwayama et al., 2019; Posfai et al., 2017; Wagner et al., 2018). Nevertheless, embryos overcome such fluctuations and achieve robust development. Mouse embryos are not prepatterned but robustly self-organize into a cyst-like structure called a blastocyst through interactions among their constituent cells. Thus, preimplantation mouse embryos have been used to determine how embryos overcome fluctuations (see reviews by Plusa and Piliszek, 2020; Saiz and Hadjantonakis, 2020; White et al., 2018; Zhang and Hiiragi, 2018; Zhu and Zernicka-Goetz, 2020). By embryonic day (E) 4.5, the late blastocyst comprises a trophectoderm (TE), an outer epithelial sheet, and an inner cell mass (ICM), which consists of a pluripotent epiblast (EPI) and a primitive endoderm (PrE). Whereas the TE and PrE cells later contribute to extraembryonic tissues, such as the placenta and yolk sac, respectively, pluripotent EPI cells give rise to all embryonic tissues, including germ cells. The EPI and PrE are formed through differentiation of the ICM during the early blastocyst stage at E3.5.

During the formation of the EPI and PrE from the early ICM, cell differentiation timing is variable. A homeodomain transcription factor, NANOG, and a zinc-finger transcription factor, GATA6, play important roles in the formation of the EPI and PrE, respectively. At the late blastocyst stage, NANOG is specifically expressed in the EPI, and is essential for its formation (Mitsui et al., 2003; Silva et al., 2009). NANOG triggers EPI specification through the coordinated expression of pluripotent transcription factors such as SOX2, OCT4, and KLF4 (Allegre et al., 2022). GATA6 is specifically expressed in the PrE, and is required for its formation (Schrode et al., 2014). At the early blastocyst stage, ICM cells co-express NANOG and GATA6. The expression of these transcription factors gradually segregates through mutual inhibition, which includes intercellular communication mediated by the FGF4–receptor tyrosine kinase (RTK)–mitogen-activated protein kinase (MAPK)–extracellular signal-regulated protein kinase (ERK) signaling pathway, resulting in NANOG single-positive and GATA6 single-positive cells by the late blastocyst stage (Bessonnard et al., 2014; Chazaud et al., 2006; Frankenberg et al., 2011; Guo et al., 2010; Kang et al., 2013; Krawchuk et al., 2013; Kurimoto et al., 2006; Messerschmidt and Kemler, 2010; Nichols et al., 2009; Ohnishi et al., 2014; Plusa et al., 2008; Yamanaka et al., 2010). NANOG and GATA6 double-positive cells are bipotential progenitor cells, and NANOG and GATA6 single-positive cells are specified to EPI and PrE fates, respectively (Saiz et al., 2016b). EPI fates are stable once specified, whereas PrE-specified cells can change their fates to EPI in rare cases (Xenopoulos et al., 2015). This fate specification takes place asynchronously throughout the blastocyst stage, and such incremental lineage allocation ensures a consistent ICM lineage composition (Saiz et al., 2016b).

In parallel with regulation by NANOG–GATA6, our recent study revealed that TEA domain transcription factor TEAD and its coactivator protein YAP also play a role in EPI formation (Hashimoto and Sasaki, 2019). TEAD and YAP are transcription factors and coactivators of the Hippo signaling pathway (Ota and Sasaki, 2008; Zhao et al., 2007; Zhao et al., 2008). The mouse genome contains four *Tead* genes (*Tead1*, *Tead2*, *Tead3*, and *Tead4*), and two *Yap*-related genes (*Yap1* and *Wwtr1* encoding the TAZ protein) (Jacquemin et al., 1996; Jacquemin et al., 1998; Kaneko and DePamphilis, 1998; Mahoney et al., 2005; Vassilev et al., 2001). In the present manuscript, we describe these four TEAD proteins and two YAP-related proteins as TEAD and YAP, respectively. Hippo signaling is activated and YAP localizes in the cytoplasm of the ICM at the early blastocyst stage (Nishioka et al., 2009). Thus, TEAD is inactive. Subsequently, YAP gradually accumulates in the nuclei, and at the late blastocyst stage, localizes in the nuclei of the EPI cells, indicating that TEAD is gradually activated during EPI formation (Hashimoto and Sasaki, 2019). A reduction in the transcriptional activity of TEAD (hereafter referred to as TEAD activity) during this process results in the reduced expression of NANOG and SOX2 (Hashimoto and Sasaki, 2019). The level of nuclear YAP, i.e., TEAD activity, at the mid-blastocyst stage is highly variable among cells, and cells with low TEAD activity are competitively eliminated via interaction with cells having high TEAD activity (Hashimoto and Sasaki, 2019). However, because both the segregation of NANOG and GATA6, and the nuclear accumulation of YAP take place asynchronously with similar timing during the blastocyst stage, it remains unknown whether TEAD–YAP plays any role in cell fate specification by NANOG–GATA6, or whether these two systems independently regulate EPI formation.

In the present study, we examined the relationship between the TEAD–YAP and NANOG–GATA6 systems during EPI formation. Detailed quantification of NANOG, GATA6, and nuclear YAP signals at the single-cell level revealed that the nuclear accumulation of YAP occurs after the segregation of NANOG and GATA6. Manipulation of these factors revealed a positive feedback loop between NANOG and TEAD activity in EPI-specified cells. The effects of TEAD–YAP on target genes also changed during the blastocyst stage, suggesting that the segregation of NANOG and GATA6 alters the transcriptional network, activates TEAD–YAP, and triggers a positive feedback loop with NANOG. The coordinated occurrence of these processes in individual cells likely ensures proper EPI formation under the condition of asynchronous EPI-fate specification.

## Results

### TEAD–YAP is activated after the segregation of NANOG and GATA6

To clarify the relationship between TEAD activity and the segregation of NANOG and GATA6, we first analyzed the expression of NANOG and/or GATA6, and the nuclear localization of YAP in freshly collected normal embryos at E3.5-E4.5. We performed immunofluorescence staining on embryos (N = 22 embryos, n = 496 cells) (Fig. 1A) and quantified nuclear signals using the single-cell resolution image analysis pipeline developed previously (Saiz et al., 2016a; Saiz et al., 2016b) (see Materials and Methods for details). Based on the expression of NANOG and/or GATA6, the cells were classified into four populations, as previously described (Saiz et al., 2016b): double-positive ICM progenitor cells (DP); NANOG single-positive EPI-specified cells (N-SP); GATA6 single-positive PrE-specified cells (G-SP); and double-negative mature EPI cells (DN) (Saiz et al., 2016a; Saiz et al., 2016b) (Fig. S1A, B). The analyzed embryos were grouped into five developmental stages according to their total cell numbers: 32–64 cells (N = 5 embryos, n = 66 cells); 64–90 cells (N = 5 embryos, n = 102 cells); 90–120 cells (N = 5 embryos, n = 116 cells); 120–150 cells (N = 3 embryos, n = 73 cells); and > 150 cells (N = 4 embryos, n = 138 cells). The 32–64 cell group corresponded to the early blastocyst stage, the 64–90 cells and 90–120 cells groups corresponded to the mid-blastocyst stage, and the 120–150 cells and >150 cells groups corresponded to the late blastocyst stage (Fig. 1A). The segregation of NANOG and GATA6 in the ICM cells was completed by the 120–150 cells stage, as reported previously (Saiz et al., 2016b) (Fig. 1B). No DN cells were present in this study (Fig. 1B).

**Figure 1.**
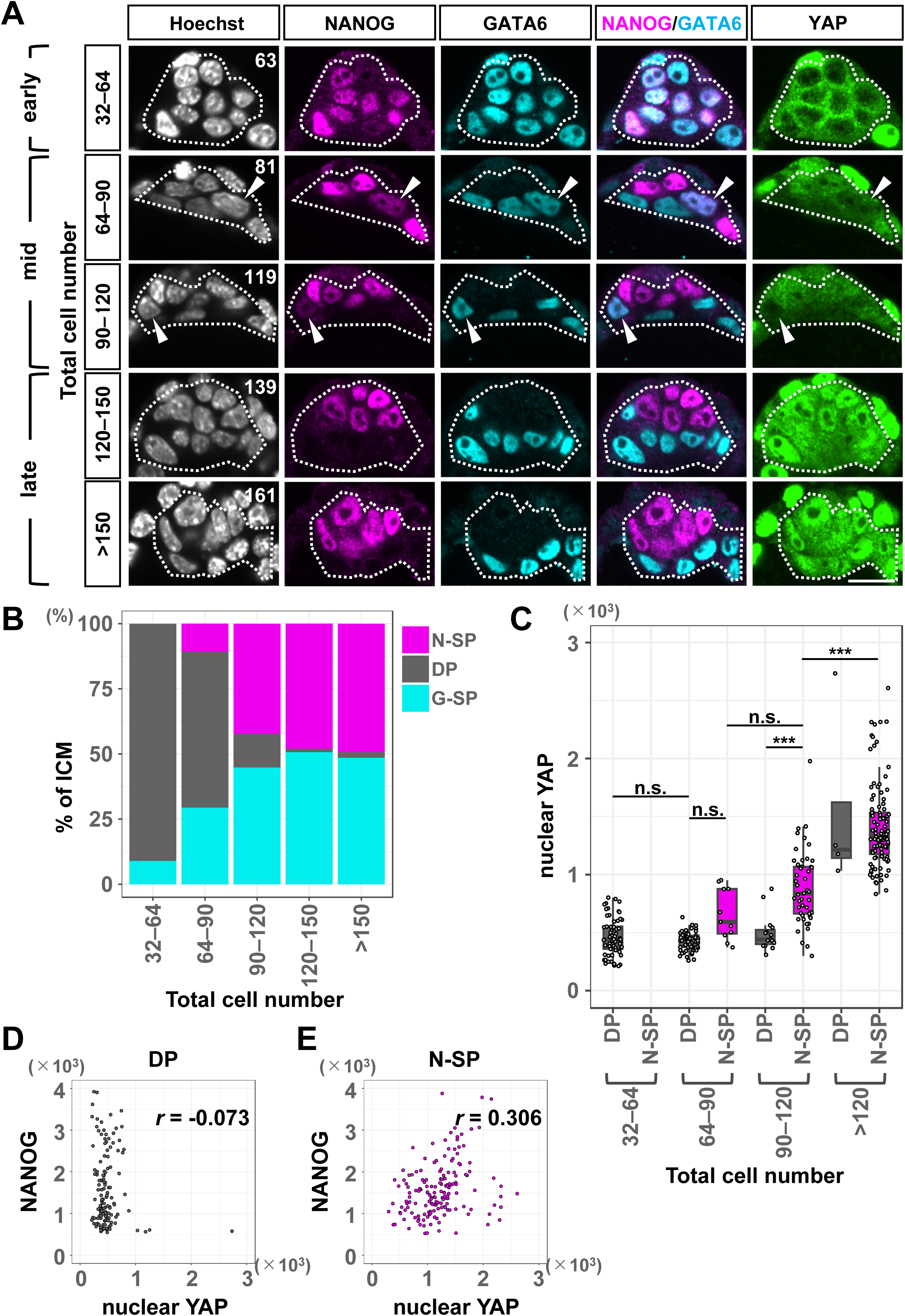
TEAD is activated after the segregation of NANOG and GATA6. (A) Representative immunofluorescence images of the ICM region of blastocysts collected at sequential developmental stages. Hoechst (nuclei), NANOG, GATA6, and YAP are shown in grayscale, magenta, cyan, and green, respectively. The total number of cells in the embryo is indicated in the Hoechst image. Dashed lines indicate the ICM. Arrowheads mark DP cells co-expressing NANOG and GATA6. The scale bar represents 20 µm. (B) Developmental changes in the average lineage composition of the ICM. The embryos were binned in the five developmental stages based on the total cell number. Magenta, gray, and cyan represent % of N-SP, DP, and G-SP cells in the ICM, respectively. The number of cells analyzed for each stage was as follows: 32–64 cells (n = 67), 64–90 cells (n = 102), 90–120 cells (n = 116), 120–150 cells (n = 73), and >150 cells (n = 138). (C) Box plots showing changes in nuclear YAP signal levels in the DP and N-SP cells during development. The embryos were binned in four developmental stages, 32–64 cells, 64–90 cells, 90–120 cells, and >120 cells. Each dot represents the nuclear YAP signal level of a cell. Sample number analyzed for each category was as follows: 32–64 cells, DP (n = 61), N-SP (n = 0); 64–90 cells, DP (n = 61), N-SP (n = 11); 90–120 cells, DP (n = 15), N-SP (n = 49), >120 cells, DP (n = 4), and N-SP (n = 103). *p*-values were determined by two-way ANOVA followed by a Tukey’s post hoc multiple comparison test. \*\*\**p* < 0.001. (D, E) Dot plots showing the correlation between NANOG and nuclear YAP signals. (F) DP cells (n = 141). (E) N-SP cells (n = 163). The correlation coefficients (*r*-values) were calculated using the Pearson correlation coefficient.

TEAD activity is regulated by the subcellular localization of YAP (Ota and Sasaki, 2008; Zhao et al., 2008). Thus, the intensity of the nuclear YAP signal was used as a proxy for TEAD activity. We compared the nuclear YAP signals of the DP and N-SP cells. DP cells in the >120 cells groups were excluded from the statistical analysis owing to the small sample size (n = 4 cells). The nuclear YAP signal remained low in the DP cells during development (Fig. 1A arrowheads, C, S1C). The nuclear YAP signal did not correlate with NANOG in the DP cells (Fig. 1A, D). The nuclear YAP signals appeared to be stronger in the N-SP cells than in the DP cells at the 64–90 cells stage, although the difference was not statistically significant. At the 90–120 cells stage, the nuclear YAP signals further increased in intensity as development progressed (Fig. 1A, C). The nuclear YAP signals exhibited a weak positive correlation with the NANOG signals in the N-SP cells (Fig. 1A, E), although the average NANOG signal levels in the N-SP cells did not change during the blastocyst stages (Fig. S1E). Nuclear YAP was also observed in the G-SP cells, but its localization occurred later than in the N-SP cells (Fig. 1A, S1C, D). Taken together, these results suggest that TEAD is activated after the segregation of NANOG and GATA6 during differentiation into the EPI.

### Segregation of NANOG and GATA6 is independent of TEAD activity

Because TEAD is activated after the segregation of NANOG and GATA6 expression, we hypothesized that the segregation of NANOG and GATA6 is independent of TEAD activity. To test this hypothesis, we generated transgenic mice that express LATS2 kinase, a negative regulator of YAP (Zhao et al., 2007), under the control of a Tet-On system. We first established embryonic stem (ES) cell lines (*R26-rtTA/+; Tg (TRE-Lats2/Egfp*)) that express reverse tetracycline trans-activator (rtTA^M2^) from the *ROSA26* locus (Hochedlinger et al., 2005), and also harbor randomly integrated multiple copies of the Tet-ON reporter cassette, which bidirectionally expresses *Lats2* and *Egfp* under the control of a tetracycline-responsive element (TRE) (Sato et al., 2007) (see Materials and Methods for details) (Fig. 2A). *R26-rtTA/+; Tg (TRE-Lats2/Egfp)* transgenic mice were produced by injecting the ES cells into 8-cell stage embryos, which resulted in the production of 100% ES cell-derived mice. Crosses of the resulting male mice and BDF1 females produced *R26-rtTA/+; Tg (TRE-Lats2/Egfp)* and *Tg (TRE-Lats2/Egfp)* embryos. The former embryos express LATS2 and EGFP upon doxycycline (Dox) treatment, whereas the latter embryos serve as negative controls that do not express LATS2 or EGFP.

**Figure 2.**
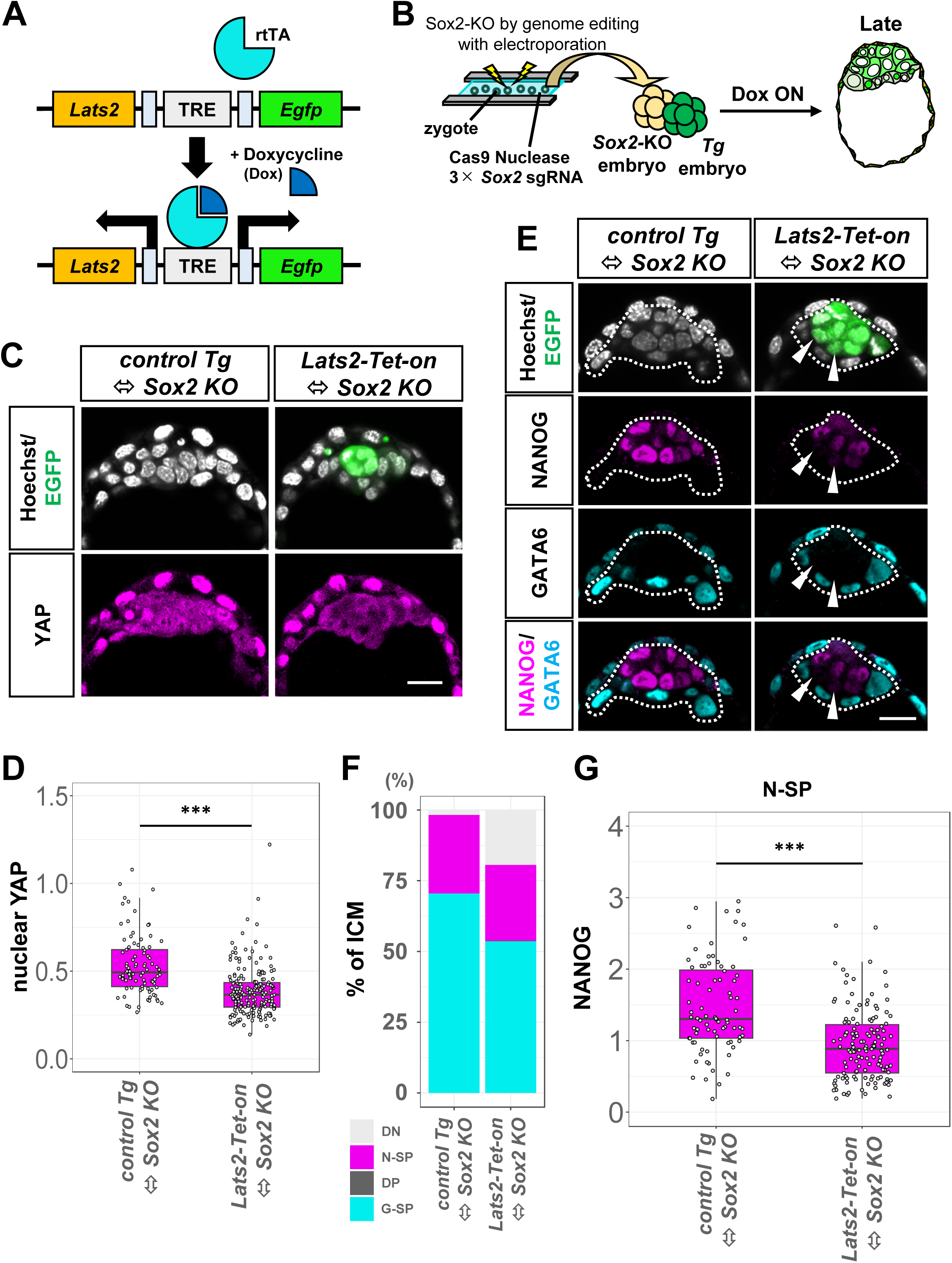
NANOG and GATA6 segregate independent of TEAD activity. (A) Scheme showing the bidirectional Tet-On system. In the presence of Dox, rtTA binds to the tetracycline responsive elements (TRE) and bidirectionally activates *Lats2* and *Egfp*. (B) Schematic of the experiments. A *Sox2-KO* (Crispant) embryo produced by genome editing with electroporation and a *Lats2-Tet-on* (or a *control Tg*) embryo were aggregated at the 8-cell stage. Chimeric embryos were cultured in Dox-containing medium and analyzed at the late blastocyst stage. (C) Representative immunofluorescence images of YAP expression in *control Tg* ⬄ *Sox2 KO* and *Lats2-Tet-on* ⬄ *Sox2 KO* embryos. Hoechst and EGFP signals are shown in white and green, respectively. EGFP was detected in the EPI area. In the *Lats2-Tet-on* ⬄ *Sox2 KO* embryos, YAP was mostly cytoplasmic in the EGFP-positive EPI cells, indicating low TEAD activity. The scale bar represents 20 µm. (D) Box plots showing nuclear YAP signal levels in SOX2-positive EPI cells of embryos shown in (C). Each dot represents the nuclear YAP signal level of a cell. The numbers of analyzed cells were as follows: *control Tg* ⬄ *Sox2 KO* (n = 89) and *Lats2-Tet-on* ⬄ *Sox2 KO* (n = 192) embryos. Student’s *t*-test; \*\*\**p* < 0.001. (E) Representative immunofluorescence images of NANOG and GATA6 expression in the *control Tg* ⬄ *Sox2 KO* and *Lats2-Tet-on* ⬄ *Sox2 KO* embryos. The expression of NANOG and GATA6 was segregated in the *Lats2-Tet-on* ⬄ *Sox2 KO* embryos. The white dashed line indicates ICM. The white arrowheads indicate DN cells. The scale bar represents 20 µm. (F) Average lineage composition of the ICM in embryos shown in (E). Light gray, magenta, dark gray, and cyan represent DN, N-SP, DP, and G-SP cells, respectively. The total numbers of analyzed cells were as follows: *control Tg* ⬄ *Sox2 KO* (n = 288), *Lats2-Tet-on* ⬄ *Sox2 KO* (n = 500). (G) Box plots showing NANOG signal levels of the N-SP cells in the embryos shown in (E). Each dot represents the NANOG signal level of a cell. The total numbers of analyzed cells were as follows: *control Tg* ⬄ *Sox2 KO* (n = 80), *Lats2-Tet-on* ⬄ *Sox2 KO* (n = 134). Student’s *t*-test; \*\*\**p* < 0.001.

We first validated the induction of LATS2 by Dox treatment. The embryos were cultured in the presence of Dox from the 8-cell stage until the early blastocyst stage. Because differentiation of TE depends on TEAD4–YAP activity (Nishioka et al., 2009; Nishioka et al., 2008; Yagi et al., 2007), activation of the Hippo signaling pathway should suppress TE development. *Tg (TRE-Lats2/Egfp)* embryos (hereafter referred to as “*control Tg* embryos”) did not express EGFP and formed blastocysts (Fig. S2A). The outer cells produced strong nuclear YAP signals and strongly expressed a TE-specific transcription factor, CDX2 (Strumpf et al., 2005). In contrast, the *R26-rtTA/+; Tg (TRE-Lats2/Egfp)* embryos (hereafter referred to as “*Lats2-Tet-on* embryos”) expressed EGFP and failed to form a TE (Fig. S2A). There was cytoplasmic localization of YAP and no CDX2 expression in all the cells, indicating that the expression of LATS2 was strong enough to activate Hippo signaling and inhibit nuclear localization of YAP. However, the absence of a TE in the *Lats2-Tet-on* embryos hindered their use for analysis of the role of TEAD–YAP during EPI formation from ICM progenitors. To overcome this problem, a *Lats2-Tet-on* embryo was aggregated at the 8-cell stage with a *Sox2*-disrupted embryo, which was generated by genome editing a zygote using the CRISPR/Cas9 system (see Materials and Methods for details). The efficiency of *Sox2* disruption was 100% (N = 18/18 embryos), as shown by the absence of a SOX2 signal (Fig. S2B). The resulting *Lats2-Tet-on* ⬄ *Sox2 KO* chimeric embryos were cultured in the presence of Dox until the late blastocyst stage (Fig. 2B). *Control Tg* ⬄ *Sox2 KO* chimeric embryos were used as controls. Because *Sox2* mutants fail to maintain an EPI at the late blastocyst stage (Avilion et al., 2003; Wicklow et al., 2014), the epiblasts of the chimeric embryos should be derived from the *Lats2-Tet-on* or *control Tg* embryos. This was confirmed in the *Lats2-Tet-on* ⬄ *Sox2 KO* embryos by the expression of EGFP in all the SOX2-positive epiblast cells (Fig. S2C).

At the late blastocyst stage, the EPI cells of the *control Tg* ⬄ *Sox2 KO* embryos produced strong nuclear YAP signals that were similar to those produced by the wildtype embryos (Fig. 2C). In contrast, YAP was excluded from the nuclei of the EPI cells of the *Lats2-Tet-on* ⬄ *Sox2 KO* embryos, indicating that TEAD was inactive (Figs. 2C, D, S2D). Using these embryos, we determined whether TEAD activity is required for the segregation of NANOG and GATA6. In both the *control Tg* ⬄ *Sox2 KO* and *Lats2-Tet-on* ⬄ *Sox2 KO* embryos a compact ICM formed, which was sandwiched between sheets of TE and PrE cells, and there were no DP cells (Fig. 2E). This indicates that the segregation of NANOG and GATA6 does not require TEAD activity. The ratio of N-SP cells to ICM cells was lower in the *control Tg* ⬄ *Sox2 KO* embryos (27%) than in the wildtype embryos (approximately 50%) (Fig. 1B, 2F). This was probably because *Sox2* mutant EPI/N-SP cells were not maintained at the late blastocyst stage (Avilion et al., 2003; Wicklow et al., 2014), and were eliminated from the forming epiblast after EPI fate specification.

### NANOG and TEAD–YAP constitute a positive feedback loop

In the N-SP cells, NANOG expression weakly correlated with the nuclear YAP signals (Fig. 1E). Consistent with this observation, in the *Lats2-Tet-on* ⬄ *Sox2 KO* embryos, in which nuclear localization of YAP was suppressed, NANOG expression in the ICM was significantly weaker than in the *control Tg* ⬄ *Sox2 KO* embryos (Fig. 2E, G). These results suggest that TEAD activity is required for the maintenance of NANOG expression in N-SP EPI-specified cells. The increase in the number of DN cells within the EPI area in the *Lats2-Tet-on* ⬄ *Sox2 KO* embryos also indicates the necessity of TEAD activity for maintaining NANOG expression (Fig. 2E arrowheads, F). The reduction in the G-SP/PrE cell ratio in the *Lats2-Tet-on* ⬄ *Sox2 KO* embryos compared to that of the *control Tg* ⬄ *Sox2 KO* embryos was likely due to the weaker expression of FGF4 in the EPI cells with reduced NANOG expression.

We next investigated whether NANOG expression also affects TEAD activity. Using a similar strategy to that employed for the overexpression of LATS2, we established a compound transgenic mouse *R26-rtTA/+; Tg (TRE-Nanog/Egfp)* that overexpressed NANOG and EGFP under the control of a Tet-ON system (Fig. 3A). BDF1 females were crossed with *R26-rtTA/+; Tg (TRE-Nanog/Egfp)* males, and the resulting embryos were treated with Dox from the 8-cell stage to the mid-blastocyst stage (Fig. 3B). The *Tg (TRE-Nanog/Egfp)* embryos (hereafter referred to as “*control Tg* embryos”) were EGFP-negative and served as controls. The EGFP-positive *R26-rtTA/+; Tg (TRE-Nanog/Egfp)* embryos (hereafter referred to as “*Nanog-Tet-on* embryos”) exhibited elevated NANOG expression in both the ICM and TE (Fig. 3C, D). There was more YAP localization in the nuclei of the ICM cells of the *Nanog-Tet-on* embryos than in the cells of the *control Tg* embryos, and there was weak correlation between the nuclear YAP signals and NANOG expression (Fig. 3C, E, F). These results suggest that NANOG promotes the nuclear localization of YAP in the ICM. Since NANOG increases TEAD activity, which is required for the maintenance of NANOG expression in EPI-specified cells, NANOG and TEAD–YAP form a positive feedback loop in EPI-specified cells, enhancing TEAD activity and maintaining NANOG expression.

**Figure 3.**
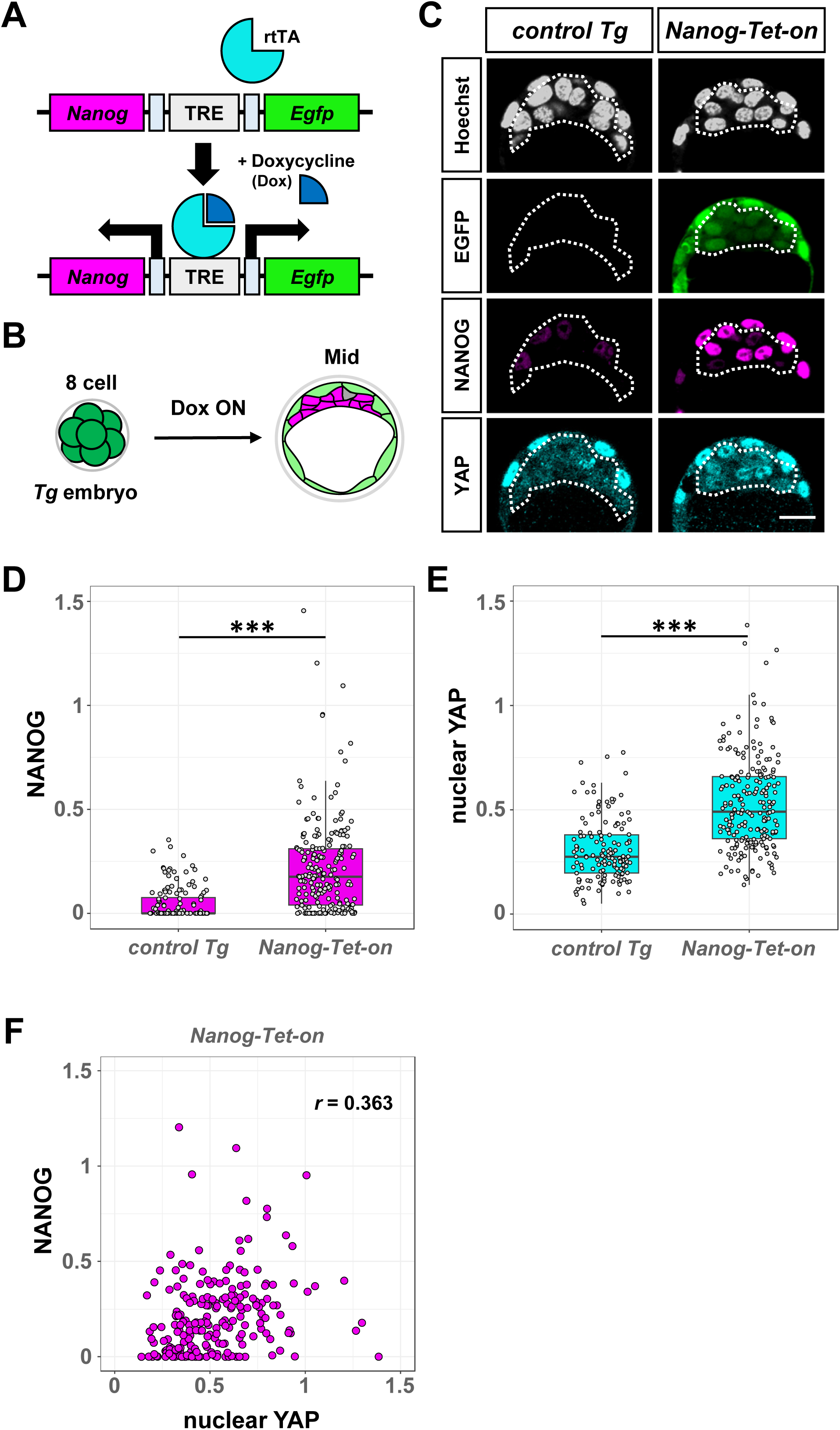
NANOG promotes nuclear localization of YAP in the ICM. (A) Scheme of the bidirectional Tet-On system. In the presence of Dox, rtTA binds to TRE and bidirectionally activates *Nanog* and *Egfp*. (B) Schematic of the experiments. The transgenic embryos were cultured in Dox-containing medium from the 8-cell stage, and analyzed at the mid-blastocyst stage. (C) Representative immunofluorescence images of NANOG and YAP expression in the *control Tg* and *Nanog-Tet-on* embryos. The white dashed line indicates the ICM. The scale bar represents 20 µm. (D, E) Box plots showing NANOG (D) and nuclear YAP (E) signal levels in the ICM of the embryos shown in (C). Each dot represents the NANOG or nuclear YAP signal level of a cell. The number of the cells analyzed in each category was as follows: *control Tg* (n = 139) and *Nanog-Tet-on* (n = 211). Student’s *t*-test; \*\*\**p* < 0.001. (F) Dot plot showing the correlation between NANOG and nuclear YAP signal levels in the *Nanog-Tet-on* embryos shown in (C) (n = 211). The correlation coefficient (*r*-value) was calculated using the Pearson correlation coefficient.

### TEAD activity is required for SOX2 upregulation in EPI-specified cells

NANOG, SOX2, and OCT3/4 constitute a core transcriptional network of naïve pluripotency in ES cells (Li and Belmonte, 2017; Macarthur et al., 2009; Ng and Surani, 2011; Orkin et al., 2008; Young, 2011). However, the expression pattern of SOX2 in preimplantation embryos differs from that of NANOG. SOX2 is initially expressed in the inner cells at the morula and early blastocyst stages and later becomes restricted to the EPI at the late blastocyst stage (Guo et al., 2010; Wicklow et al., 2014). To explore the relationship between EPI fate specification and SOX2 expression, we examined SOX2 expression in various cell types categorized by NANOG and GATA6 expression. Embryos were immunofluorescently stained for SOX2, NANOG, and GATA6, and the signals were quantified as described above (N = 24 embryos, n = 542 cells) (Fig. 4A). SOX2 expression remained low in the DP cells, but gradually increased in N-SP cells after the 90-cell stage (Fig. 4A, B). Conversely, SOX2 expression gradually decreased in G-SP cells as development progressed (Fig. 4A, S3A). Thus, SOX2 expression was upregulated following the segregation of NANOG and GATA6.

**Figure 4.**
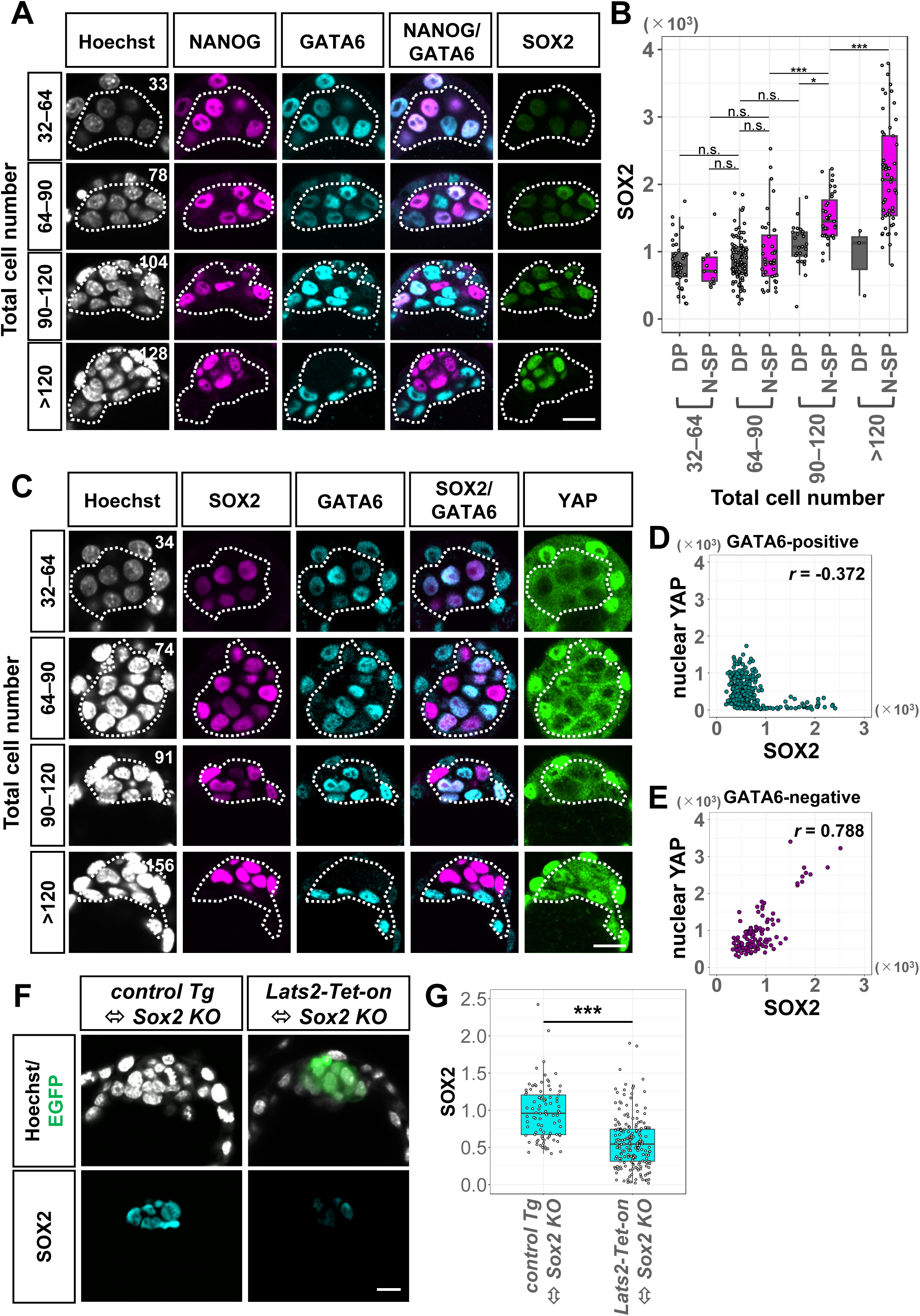
TEAD activity is required for the upregulation of SOX2 in EPI-specified cells. (A) Representative immunofluorescence images of NANOG, GATA6, and SOX2 expression in the ICM of the blastocyst-stage embryos. Hoechst (nuclei), NANOG, GATA6, and SOX2 are shown in grayscale, magenta, cyan, and green, respectively. Embryos were categorized into four stages depending on the total cell number. The total number of cells in the embryo is indicated in the Hoechst image. The dashed lines indicate the ICM. The scale bar represents 20 µm. (B) Box plots showing SOX2 signal levels in the DP and N-SP cells of the embryos shown in (A). Each dot represents the SOX2 signal level of a cell. The cell numbers analyzed in each category were as follows: 32–64 cells, DP (n = 40), N-SP (n = 11); 64–90 cells, DP (n = 92), N-SP (n = 40); 90–120 cells, DP (n = 25), N-SP (n = 35), >120, DP (n = 3), N-SP (n = 53). Two-way ANOVA followed by a Tukey’s post hoc multiple comparison test. \**p* < 0.05, \*\*\**p* < 0.001, n.s. = not significant. (C) Representative immunofluorescence images of SOX2, GATA6, and YAP expression in the ICM of the blastocyst-stage embryos. Hoechst (nuclei), SOX2, GATA6, and YAP are shown in grayscale, magenta, cyan, and green, respectively. The embryos were categorized into four stages depending on the total cell number. The total number of cells in the embryo is indicated in the Hoechst image. The dashed lines indicate the ICM. The scale bar represents 20 µm. (D, E) Dot plots showing correlations between SOX2 and nuclear YAP signal levels in the GATA6-positive (n = 322) (D) and GATA6-negative (n = 111) (E) cells. The correlation coefficient (*r*-value) was calculated using the Pearson correlation coefficient. (F) Representative immunofluorescence images of SOX2 expression in the *control Tg* ⬄ *Sox2 KO* and *Lats2-Tet-on* ⬄ *Sox2 KO* embryos. Hoechst and EGFP signals are shown in grayscale and green, respectively. The scale bar represents 20 µm. (G) Box plots showing SOX2 signal levels in SOX2-positive cells of the embryos shown in (F). Each dot represents the SOX2 signal level of a cell. The numbers of analyzed cells were as follows: *control Tg* ⬄ *Sox2 KO* (n = 89) and *Lats2-Tet-on* ⬄ *Sox2 KO* embryos (n = 192). Student’s *t*-test; \*\*\**p* < 0.001.

Given that increase in SOX2 expression in N-SP cells resembled the increase in nuclear YAP in these cells (Fig. 1C, 4B), we next examined the relationship between SOX2 expression and nuclear YAP. Using immunofluorescence for YAP, SOX2, and GATA6, we used GATA6-negative ICM cells as a proxy for N-SP cells (N = 23 embryos, n = 433 cells) (Fig. 4C, S3B, C). In GATA6-positive cells representing DP and G-SP cells, SOX2 expression did not correlate with nuclear YAP (Fig. 4C, D). However, in GATA6-negative cells, there was a positive correlation between SOX2 expression and nuclear YAP (r = 0.788) (Fig. 4C, E).

To assess the involvement of TEAD activity in SOX2 expression in N-SP cells, we examined SOX2 expression in *Lats2-Tet-on* ⬄ *Sox2 KO* embryos, where LATS2 overexpression inhibited nuclear localization of YAP in EPI cells, at the late blastocyst stage (Fig. 2B). SOX2 expression was significantly lower in *Lats2-Tet-on* ⬄ *Sox2 KO* embryos compared to *control Tg* ⬄ *Sox2 KO* chimeric embryos (Fig. 4F, G). These results imply that TEAD activity is essential for strong SOX2 expression at the late blastocyst stage, consistent with previous findings (Hashimoto and Sasaki, 2019). Therefore, SOX2 upregulation in EPI-specified cells depends on TEAD activity.

### Effects of TEAD–YAP on *Nanog* and *Sox2* are different

We demonstrated that TEAD activity is required for the strong expression of both NANOG and SOX2 at the late blastocyst stage. Next, we examined whether TEAD activation is sufficient to induce these transcription factors. To promote nuclear localization of YAP, blastocyst stage embryos were treated with the LATS inhibitor (LATSi, also known as TRULI) (Kastan et al., 2021), which induces clear nuclear localization of YAP in the ICM within 2 hours (Otsuka et al., 2023). To investigate the effects of the timing of TEAD–YAP activation on gene expression, embryos were treated with LATSi for 12 hours at 0–12, 12–24, or 24–36 hours after blastocoel formation (Fig. 5A).

**Figure 5.**
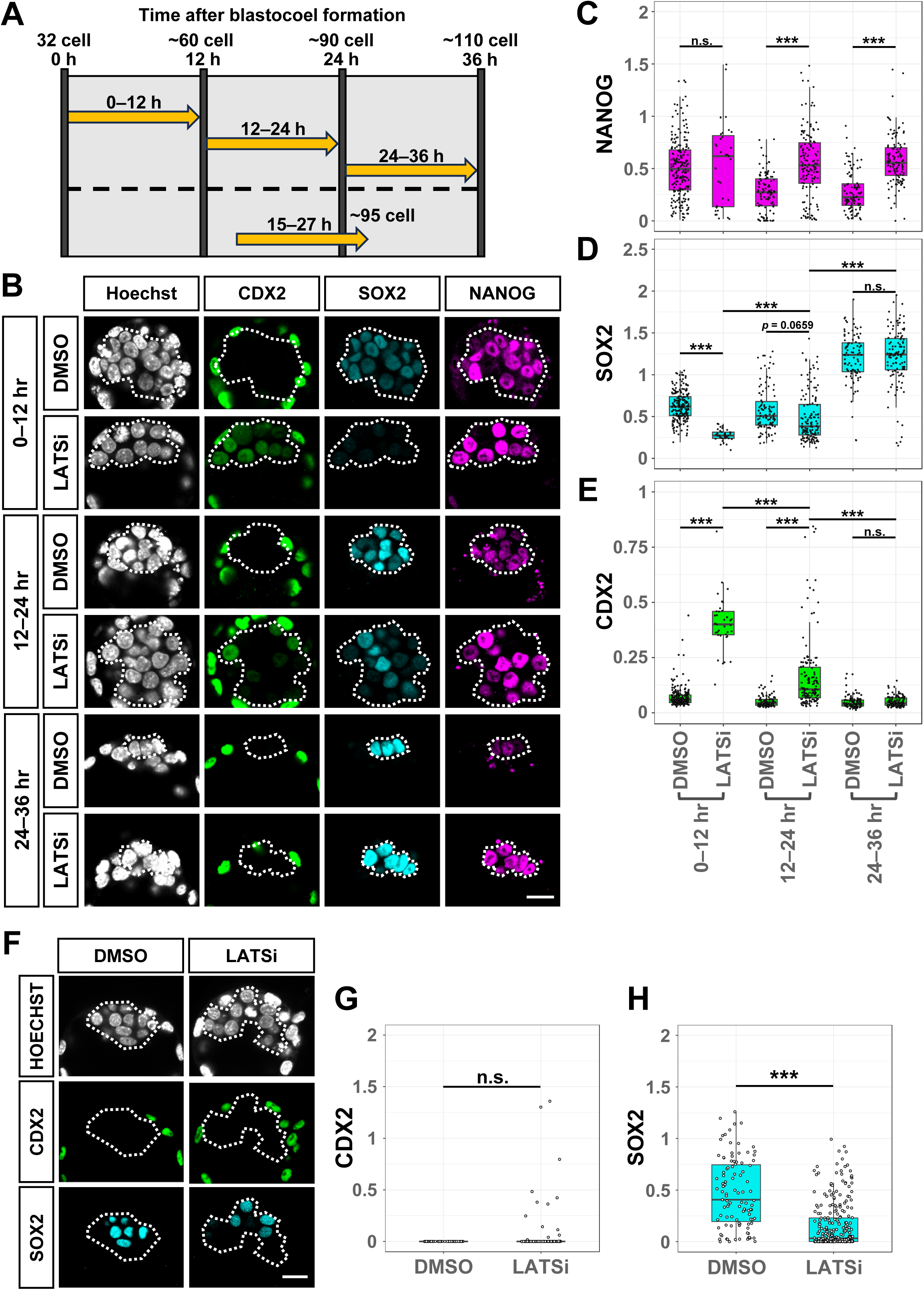
Responses of *Cdx2* and *Sox2* to TEAD–YAP change during the blastocyst stage. (A) Schematic of the experiments. Embryos were cultured in a medium containing a LATS inhibitor (LATSi) for 12 h starting from 0, 12, 15, or 24 h after blastocoel formation, and were subsequently analyzed. (B) Representative immunofluorescence images of expression of CDX2, SOX2, and NANOG in embryos treated with LATS inhibitor for 12 h starting from 0, 12, or 24 h after blastocoel formation. The dashed lines indicate the ICM. The scale bar represents 20 µm. (C, D, E) Box plots of NANOG (C), SOX2 (D), and CDX2(E) signal levels of ICM cells of embryos shown in (B). Each dot represents the signal level of a cell. The numbers of analyzed cells in each category were as follows: 0–12 h, DMSO (n = 210), LATSi (n = 35); 12–24 h, DMSO (n = 103), LATSi (n = 134); 24–36 h, DMSO (n = 96), LATSi (n = 115). Two-way ANOVA followed by a Tukey’s post hoc multiple comparison test. \*\*\**p* < 0.001, n.s. = not significant. (F) Representative immunofluorescence images of the expression of CDX2 and SOX2 in the embryos treated with LATSi for 15–27 h. The dashed lines indicate the ICM. The scale bar represents 20 µm. (G, H) Box plots of CDX2 (G) and SOX2 (H) signal levels in the ICM of the embryos shown in (F). Each dot represents the signal level of a cell. The numbers of analyzed cells were as follows: DMSO (n = 103) and LATSi (n = 277). Student’s *t*-test; \*\*\**p* < 0.001.

We first examined the effects on NANOG expression. LATSi treatment over 12–24 or 24–36 hours significantly increased the expression level of NANOG compared to control dimethyl sulfoxide (DMSO)-treated embryos, indicating that the activation of TEAD is sufficient for the induction of NANOG (Fig. 5B, C). These results are consistent with the hypothesis that TEAD–YAP and NANOG constitute a positive feedback loop in EPI-specified cells. We next examined the effects on SOX2 expression. LATSi treatment over 12–24 or 24–36 hours did not significantly alter the expression level of SOX2 compared to control DMSO-treated embryos (Fig. 5B, D). Therefore, the activation of TEAD is insufficient for induction of SOX2, raising the possibility that the positive effect of TEAD–YAP on the expression of SOX2 in EPI-specified cells is indirect. Taken together, these results suggest that TEAD–YAP positively regulates *Nanog*, whereas it has no direct effect on *Sox2* in EPI-specified cells.

### Response to TEAD–YAP changes with EPI fate specification

To further explore the effects of TEAD–YAP on NANOG and SOX2, we treated embryos with LATSi over 0–12 hours after blastocoel formation (Fig. 5A), when the majority of the ICM cells are DP and TEAD is inactive (Fig. 1). While LATSi treatment did not significantly alter the expression level of NANOG, it almost completely suppressed the expression of SOX2. These effects are clearly different from the effects of LATSi treatment over 24–36 hours, which represent the effects on EPI-specified cells, suggesting that the effects of TEAD–YAP change with EPI fate specification.

To further explore the differential effects of TEAD–YAP between DP and EPI-specified cells, we examined the effects on a TE-specific transcription factor, CDX2, because early ICM cells maintain the ability to express CDX2 (Nishioka et al., 2009; Posfai et al., 2017). In the ICM of the 0–12 hour LATSi-treated embryos, CDX2 was strongly expressed in all cells (Fig. 5B, E). In the 12–24 hour LATSi-treated embryos, both CDX2 and SOX2 were expressed in a non-overlapping manner (Fig. 5B, D, E). In the 24–36 hours LATSi-treated embryos, CDX2 was not expressed (Fig. 5B, E). Thus, the effects of TEAD–YAP on CDX2 also change with EPI fate specification.

The ICM of the 12–24 hour LATSi-treated embryos showed non-overlapping expression of CDX2 and SOX2, raising the possibility that the expression of CDX2 inhibited *Sox2* expression. In the ICM of the 15–27 hour LATSi-treated embryos, the majority of the ICM cells did not express either CDX2 or SOX2, and the scattered expression of CDX2 and SOX2 did not overlap (Figs. 5A, F-H, S4). This result suggests that the suppression of SOX2 is independent of CDX2. It is likely that TEAD–YAP directly suppresses *Sox2*, as demonstrated in morula-stage embryos (Frum et al., 2019). Taken together, these results suggest that the response of TEAD–YAP target genes changes with EPI-fate specification: TEAD–YAP activates *Cdx2* and represses *Sox2* before EPI fate specification, whereas it activates *Nanog* after the specification.

## Discussion

In this study, we elucidated the interplay between two transcription factor systems—NANOG–GATA6 and TEAD–YAP—that regulate EPI formation. We investigated the dynamics of these transcription factors during EPI formation in detail, and studied their relationships by genetic and pharmacological manipulation of embryos. The results, combined with previous findings, are summarized in the following model (Fig. 6).

**Figure 6.**
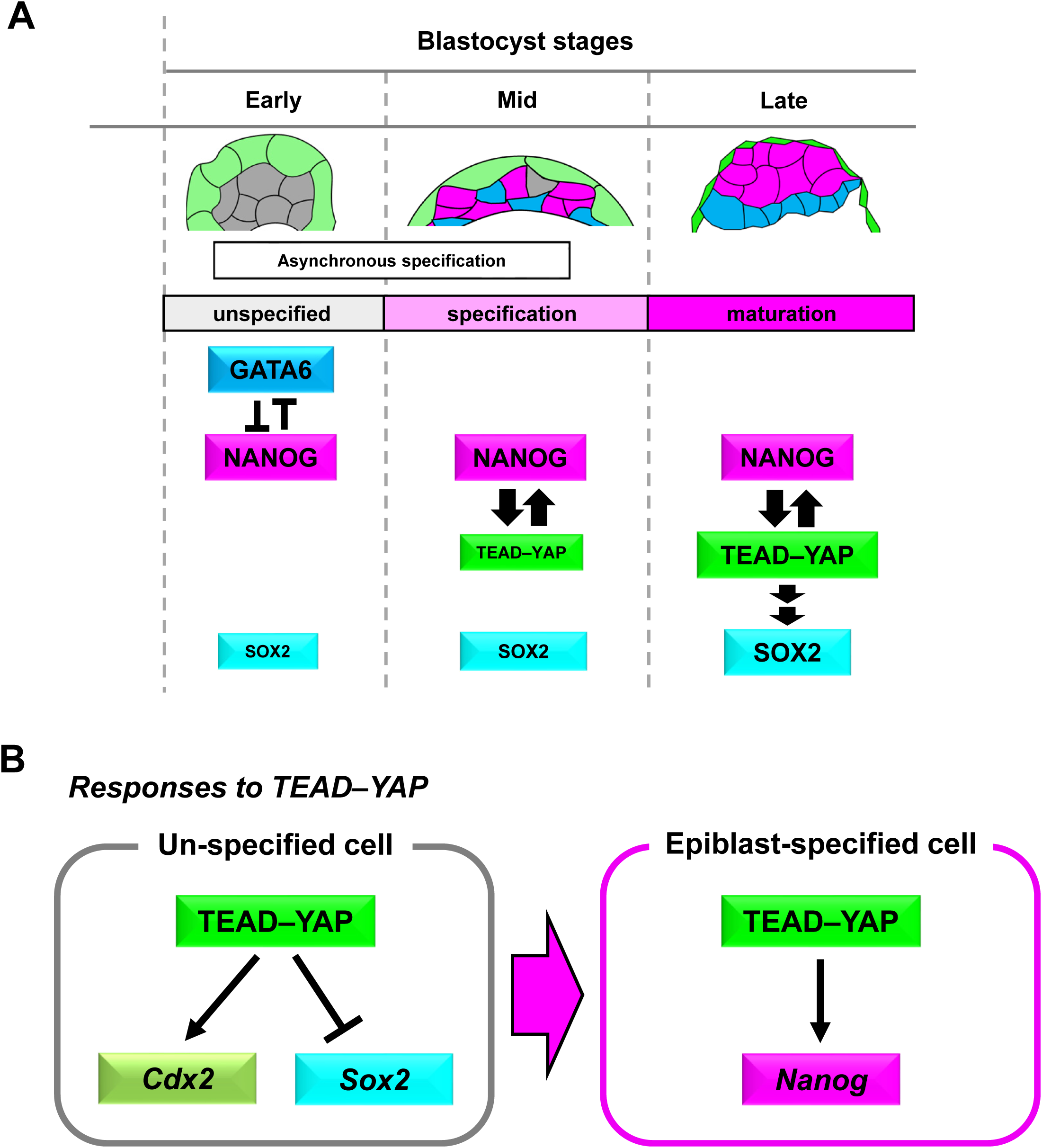
Model of transcriptional regulation of epiblast formation. (A) Model of epiblast formation during the blastocyst stage regulated by interaction of transcription factors. NANOG and GATA6 are co-expressed in unspecified ICM cells. EPI is specified by the segregation of NANOG from GATA6, which is independent of TEAD activity. Once specified, TEAD is activated in EPI cells. NANOG and TEAD activity constitutes a positive feedback loop, which increases TEAD activity and maintains NANOG expression. TEAD activity is required for upregulation of SOX2, but its effect is indirect. These transcription factors support the maturation of the epiblast. (B) Model of transcriptional network change with epiblast fate specification. In unspecified cells, TEAD–YAP activates *Cdx2* and represses *Sox2*. In EPI-specified cells, TEAD–YAP induces *Nanog*.

Early blastocyst stage ICM cells are bipotential progenitors that can differentiate into either EPI or PrE, co-expressing NANOG and GATA6 and exhibiting cytoplasmic YAP (Nishioka et al., 2009; Plusa et al., 2008). As EPI formation progresses, NANOG segregates from GATA6, and YAP translocates to the nucleus asynchronously among ICM cells (Hashimoto and Sasaki, 2019; Saiz et al., 2016b), resulting in a mixture of cells with variable gene expression. Our quantitative analysis revealed that nuclear YAP accumulation occurs in N-SP cells after the segregation of NANOG from GATA6 (Fig. 1). Consistent with this, the segregation of NANOG and GATA6 is independent of TEAD activity (Fig. 2). However, TEAD activity affects the PrE/EPI ratio. The PrE/EPI ratio is reduced in embryos with reduced TEAD activity (Fig. 2F), probably due to the reduced expression of FGF4 in EPI cells.

Immediately after segregation, the emerging N-SP cells represent the EPI-specified cells (Saiz et al., 2016b). Once specified, EPI fate becomes stabilized (Xenopoulos et al., 2015). We demonstrated that NANOG and TEAD activity form a positive feedback loop in N-SP cells: NANOG promotes nuclear YAP (increasing TEAD activity) (Fig. 3), and TEAD activity maintains NANOG expression (Figs. 2, 5). This positive feedback loop likely contributes to the fate stabilization of EPI-specified cells. TEAD–YAP activity is also necessary for SOX2 upregulation in N-SP cells (Fig. 4), although this effect is indirect as discussed below. It is likely that the sustained expression of NANOG and the upregulation of SOX2 via TEAD–YAP in N-SP cells support the transition of EPI-specified cells to mature EPI cells, which have naïve pluripotency (Nichols et al., 2009) (Fig. 6A).

TEAD activity is essential for the robust expression of both NANOG and SOX2. However, the effects of TEAD activity on NANOG and SOX2 were different. Whereas increased TEAD activity upregulated NANOG, it did not alter the SOX2 expression level (Fig. 5). This difference could originate from the differences in the modes of target gene regulation by TEAD–YAP. In ES cells, YAP1 binds 4 kb upstream of *Nanog* but not around *Sox2* (Sun et al., 2020) (Fig. S5A, B). Therefore, in EPI-specified cells, TEAD–YAP directly regulates *Nanog*, whereas the effect of TEAD–YAP on *Sox2* is indirect.

The response of ICM cells to TEAD–YAP changes during the blastocyst stage (Fig. 6B). Early blastocyst ICM cells still retain the ability to differentiate into TE (Posfai et al., 2017; Rossant and Lis, 1979; Spindle, 1978) and exhibit characteristics of morula-stage embryos, such as *Cdx2* responsiveness to TEAD–YAP (Posfai et al., 2017)(Nishioka et al., 2009). Our results indicate that TEAD–YAP suppresses *Sox2* at the early blastocyst stage (Fig. 5), demonstrating additional similarities between the early ICM and the morula. Thus, *Sox2* expression at the early blastocyst stage is supported by the absence of TEAD–YAP repression (this study) and activation by NANOG and OCT3/4 (Frum et al., 2019). These regulatory mechanisms are gradually lost as development progresses, and at the late blastocyst stage, TEAD–YAP positively regulates *Nanog*.

Most early ICM cells are DP and gradually transition to N-SP or G-SP cells. Thus, DP cells retain transcriptional regulatory mechanisms akin to the morula stage, where TEAD–YAP activates *Cdx2* and represses *Sox2*. Upon EPI fate specification in N-SP cells, TEAD–YAP’s regulatory role shifts to activating *Nanog* and ceasing *Sox2* repression. This indicates a developmental transition in TEAD–YAP target gene regulation as cells move from DP to EPI-specified N-SP cells. The segregation of NANOG and GATA6 alters the transcriptional network from morula-like to EPI-specified, triggering TEAD–YAP activation to support proper EPI development. Because cell fate specification from DP to N-SP/EPI or G-SP/PrE cells takes place asynchronously to ensure the constant EPI/PrE ratio (Saiz et al., 2016b), the coordination of changes in the transcriptional network and TEAD–YAP activation at the time of EPI fate specification in individual cells is important for the proper formation of EPI cells.

During EPI formation, differences in TEAD activity induce cell competition (Hashimoto and Sasaki, 2019). Cells with high TEAD activity and strong pluripotency factor expression become winners, while cells with low TEAD activity and weaker pluripotency factor expression are eliminated as losers through apoptosis. Our results reveal how variations in TEAD activity occur in the ICM. YAP nuclear localization takes place in EPI-specified N-SP cells (Fig. 1). Thus, asynchronous cell fate specification in ICM cells (Saiz et al., 2016b) results in varying nuclear YAP levels. Whereas EPI-specified cells with nuclear YAP become winners, progenitor DP cells with cytoplasmic YAP become losers. This is consistent with a previous observation that the loser cells weakly co-express SOX2 and a PrE-specific transcription factor, SOX17 (Hashimoto and Sasaki, 2019). However, because not all progenitor cells are eliminated through cell competition, the mechanism of loser fate specification requires further clarification.

## Materials and methods

### Mouse lines

B6D2F1/Jcl (CLEA, Tokyo, Japan) (hereafter referred to as BDF1), *Gt (ROSA) 26Sor^tm1(rtTA*M2)^ ^Jae^*(abbreviated as *R26-rtTA*) (RRID:MMRRC_014112-MU) (Hochedlinger et al., 2005), *R26-rtTA/+; Tg (TRE-Lats2/Egfp)* (this study), and *R26-rtTA/+; Tg (TRE-Nanog/Egfp)* (this study) mice were used. The mice were housed in environmentally controlled rooms at the Animal Facility of the Graduate School of Frontier Biosciences, Osaka University. All experiments with mice had the approval of the Animal Care and Use Committee of the Graduate School of Frontier Biosciences, Osaka University, and all recombinant DNA experiments had the approval of the Gene Modification Experiments Safety Committee of Osaka University.

### Embryo collection and culture

Preimplantation embryos were collected by following the standard protocol with slight modifications (Behringer et al., 2014). Briefly, BDF1 female mice superovulated by intraperitoneal injection of 10 IU pregnant mare serum gonadotropin and human chorionic gonadotropin at 46–52 h intervals were crossed with BDF1 male mice, *R26-rtTA/+; Tg (TRE-Lats2/Egfp)* male mice, or *R26-rtTA/+; Tg (TRE-Nanog/Egfp)* male mice. Zygotes at the 1-cell stage were obtained from the E0.5 oviduct ampulla and treated with hyaluronidase to remove cumulus cells. 2-cell embryos were obtained from the E1.5 oviduct. These embryos were cultured in a well of a 72-well MiniTray (Nunc 136528) containing 10 µL of KSOM-AA medium (ARK Resource, Kumamoto, Japan), covered with 5 µL of mineral oil, at 37 °C in a 5% CO_2_ incubator. Blastocysts for gene expression analyses of normal embryos were collected from the uterus of the mice by flushing with M2 medium between E3.5 and E4.5.

### Plasmids

A Tol2 transposase expression plasmid, pCAGGS-T2TP (Kawakami et al., 2004), and a bidirectional Tet-ON reporter plasmid, pT2K-BI-TRE-EGFP (Sato et al., 2007), were kindly provided by Dr. Koichi Kawakami and Dr. Yoshiko Takahashi. The coding sequence of HA-tagged mouse *Lats2* cDNA (Ota and Sasaki, 2008) and mouse *Nanog* cDNA were cloned into pT2K-BI-TRE-EGFP to create pT2K-BI-TRE-HA-Lats2/EGFP. A hyperactive piggyBac transposase expression plasmid, pCMV-hyPBase (Yusa et al., 2011) was obtained from the Wellcome Trust Sanger Institute. pPB-LR51-EF1a-puroCas9 (Koike-Yusa et al., 2014) was kindly provided by Dr. Kosuke Yusa and the Wellcome Trust Sanger Institute. A bidirectional expression plasmid, pPB-BI-TRE-Nanog/EGFP, was constructed by modifying pPB-LR51-EF1a-puroCas9. The plasmid consists of PB terminal repeats, *EGFP* cDNA, TRE 3G BI promoter of Tet-On 3G Bidirectional Inducible Expression System (EF1alpha Version), *Nanog* cDNA, and poly (A) signals. Sequence information for this plasmid is available upon request. pCAGGS-IP (Niwa et al., 2005) was kindly provided by Dr. Hitoshi Niwa. pX330-U6-Chimeric_BB-CBh-hSpCas9 (hereafter referred to as pX330) (Cong et al., 2013) was obtained from Addgene.

### Production of transgenic mice

We first established ES cell lines from blastocysts obtained by crossing B6D2F1/Jcl female and *Gt (ROSA)26Sor^tm1^ ^(rtTA*M2)^ ^Jae^* male mice. *Gt (ROSA)26Sor^tm1^ ^(rtTA*M2)^ ^Jae^* (RRID:MMRRC_014112-MU) (Hochedlinger et al., 2005) mice were obtained from MMRRC. The resulting ES cell lines were named *R26-rtTA/+* cells. One ES cell line (# 3) was used after confirming efficient chimera formation and germline transmission. *R26-rtTA/+* cells were co-transfected with pCAGGS-T2TP, pT2K-BI-TRE-HA-Lats2/EGFP, and pCAGGS-IP using Lipofectamine 2000, as described previously (Tamm et al., 2016). Transfected cells were selected with 1 μg/mL puromycin for 2 days followed by trypsinization for passage. The single cell-derived ES cell colonies were treated with 1 μg/mL DOX for 24 h, and EGFP-positive colonies were manually picked using a fluorescent microscope and expanded. After expansion, the ES cells were treated with 1 μg/mL Dox for 24 h, and the resulting clones, which were confirmed by strong EGFP expression, were selected and named *R26-rtTA/+; Tg (TRE-Lats2/Egfp)* cells; one ES line (# 2) was used to produce mice. The resulting F0 mice were named *R26-rtTA/+; Tg (TRE-Lats2/Egfp)* and were used for analysis. Using essentially the same protocol, pPB-BI-TRE-Nanog/EGFP and pCMV-hyPBase were used instead of pT2K-BI-TRE-Lats2/EGFP and pCAGGS-T2TP, respectively, to produce *R26-rtTA/+; Tg (TRE-Nanog/Egfp)* ES cells, and clone #3 was used to produce *R26-rtTA/+; Tg (TRE-Nanog/Egfp)* mice.

### Generation of chimeric embryos

To generate chimeric embryos, superovulated wildtype BDF1 females were crossed with *R26-rtTA/+; Tg (TRE-Lats2/Egfp)* males. The transgenic embryos, *R26-rtTA/+; Tg (TRE-Lats2/Egfp)* or *Tg (TRE-Lats2/Egfp)*, were recovered at the 1- or 2-cell stage, and cultured *in vitro* until the 8-cell stage. In parallel, BDF1 embryos were recovered at the 1-cell stage, and genome-edited by introducing Cas9 protein (Integrated DNA Technologies) and three *Sox2* sgRNAs by electroporation (Hashimoto et al., 2016). The template DNA fragments for the three *Sox2* sgRNAs were prepared in separate PCR reactions using one of the following *Sox2* forward primers: (5 ′-TTAATACGACTCACTATAGGGCCCGCAGCAAGCTTCGGGTTTTAGAGCTAGAAATAGCAAGTTAAAAT-3 ′, 5 ′ -TTAATACGACTCACTATAGGGCCTCAACGCTCACGGCGGTTTTAGAGCTAGAAATAGCAAGTTAAAAT-3 ′, 5 ′ -TTAATACGACTCACTATAGGGCTCTGTGGTCAAGTCCGGTTTTAGAGCTAGAAATAGCAAGTTAAAAT-3 ′); a common reverse primer (3 ′ -AAAAGCACCGACTCGGTGCCACTTTT-3′); and a template pX330 plasmid. *In vitro* transcription and purification of *Sox2* sgRNAs were performed as described in the literature (Hashimoto et al., 2016). Chimeric embryos were generated by aggregating *Sox2*-Crispant embryos with *R26-rtTA/+; Tg (TRE-Lats2/Egfp)* or *R26-rtTA/+;Tg (TRE-Lats2/Egfp)* transgenic embryos. At the 8-cell stage, zona pellucida of *Sox2*-Crispant and transgenic embryos were removed by brief incubation in acidic Tyrode’s solution. The denuded embryos were aggregated by culturing in small wells in pairs of *Sox2*-Crispant and transgenic embryos in KSOM medium supplemented with 1 μg/mL Dox until the late blastocyst stage.

### LATS inhibitor treatment

For the inhibitor treatments, embryos were cultured in KSOM medium containing 10 μM LATS inhibitor (TRULI) (MedChemExpress). The control embryos were cultured in KSOM media containing the same concentrations of DMSO.

### Immunofluorescence staining and confocal image acquisition

The embryos were stained with an immunofluorescent agent using a 72-well MiniTray (Nunc 136528), as previously described with slight modifications. Briefly, the embryos were fixed with 4% paraformaldehyde in Dulbecco’s phosphate-buffered saline (PBS) for 15 min at room temperature, then washed and permeabilized with 0.1% Triton X-100 in PBS (PBST) for 30 s twice at room temperature. The embryos were then blocked with 2% donkey serum in PBST (blocking solution) for 5 min, and incubated overnight with the primary antibodies diluted with blocking solution in a 1:100 ratio at 4 °C. We used the following primary antibodies: rabbit anti-NANOG (ReproCell, RCAB0001P), rat anti-NANOG (Invitrogen, 14-5761-80), rabbit anti-SOX2 (Cell Signaling Technology, D9B8N), goat anti-GATA6 (R&D System, AF1700), rabbit anti-YAP1 (Cell Signaling Technology, 14074S), and mouse anti-CDX2 (BioGenex, MU392-UC). For some experiments, we also used the following prelabelled antibodies: Alexa Fluor 488 goat anti-SOX2 (R&D System, AF2018), and Alexa Fluor 647 goat anti-GATA6 (R&D System, AF1700). These prelabelled antibodies were prepared in house with Molecular Probes Antibody Labeling Kits (Thermo Fisher Scientific) according to the manufacturer’s instructions. The embryos were washed twice in PBST for 1 min, then incubated with the secondary antibodies diluted with PBST at a ratio of 1:1000 and 1 μg/mL Hoechst 33342 (Dojindo, H342) for more than 1 h at room temperature. We used the following secondary antibodies: donkey anti-rabbit IgG Alexa Fluor Plus 488 (Thermo Fisher Scientific, A32790), donkey anti-rat IgG Alexa Fluor 488 (Thermo Fisher Scientific, A21208), donkey anti-mouse IgG Alexa Fluor Plus 488 (Thermo Fisher Scientific, A32766), donkey anti-rabbit IgG Alexa Fluor Plus 555 (Thermo Fisher Scientific, A32794), donkey anti-rabbit IgG Alexa Fluor 647 (Thermo Fisher Scientific, A31573), donkey anti-rat IgG Alexa Fluor Plus 647 (Thermo Fisher Scientific, A48272), donkey anti-mouse IgG Alexa Fluor Plus 647 (Thermo Fisher Scientific, A32787), and donkey anti-goat IgG Alexa Fluor 647 (Thermo Fisher Scientific, A21447). For observation, the stained embryos were washed twice with 1% bovine serum albumin in PBS, and added to a small drop of PBS on a glass-bottomed dish (IWAKI).

Confocal images were obtained using a spinning disk confocal microscope system consisting of a Nikon Ti2 microscope (Nikon Solutions Inc.), a Dragonfly system (Oxford Instruments Ltd), and a Zyla sCOMS camera (Oxford Instruments Ltd), or a Nikon A1 point-scanning confocal microscope system (Nikon Solutions Inc.). The former and latter systems were used for observation of the normal and manipulated embryos, respectively.

### Data processing and quantification of the confocal images

The fluorescent signals produced by the normal embryos were quantified in detail according to the pipeline developed by Hadjantonakis’ group (Saiz et al., 2016a; Saiz et al., 2016b). Briefly, MATLAB (ver. 2012)-based MINS software was used to segment nuclei and quantitate nuclear signals (Saiz et al., 2016a), with manual confirmation/correction of segmentation and identification of ICM–TE cells using Image J (Schneider et al., 2012). Compensation of fluorescence intensity decay along the z-axis and categorization of cell types based on NANOG and GATA6 signal intensities with k-means clustering were performed using the R scripts described in the literature (Saiz et al., 2016b).

We quantified the fluorescence signals produced by the manipulated embryos using IMARIS software (Oxford Instruments Ltd), as described previously (Hashimoto and Sasaki, 2019). Briefly, we quantified nuclear fluorescence signals in regions of interest comprising 6 μm-diameter spheres. The fluorescence signals were divided by the Hoechst signals to compensate for signal decay along the z-axis.

### Statistical analysis

Statistical analysis was performed using Prism 9 software (GraphPad Software, MA, USA). The statistical methods and the numbers of samples are described in the figure legends.

## Acknowledgments

We thank Dr. Anna-Katerina Hadjantonakis and Dr. Sayali Chowdhary for the MINS program and advice on its use; Dr. Shin-ichi Kawaguchi and Dr. Ritsuko Suyama for advice on R; Dr. Koichi Kawakami and Dr. Yoshiko Takahashi for the pCAGGS-T2TP and pT2K-BI-TRE-EGFP plasmids; Dr. Hitoshi Niwa for the pCAGGS-IP plasmid and Nanog cDNA; Dr. Kosuke Yusa and the Wellcome Trust Sanger Institute for the pPB-LR51-EF1a-puroCas9 plasmid; Dr. Nancy L. Craig and the Wellcome Trust Sanger Institute for the pCMV-hyPBase plasmid; Dr. Feng Zhang and Addgene for the pX330 plasmid; and Dr. Rudorf Jaenish and MMRRC for the *Gt(ROSA)26Sor^tm1(rtTA*M2)Jae^*mice.

## Footnotes

## Author contributions

Conceptualization: H.Sasaki., N.H.; Methodology: H. Sasaki, N.H., M.H., H.M., H. Shimojo.; Software: H. Sasaki, N.H.; Validation: H. Sasaki, N.H.; Formal analysis: N.H., M.H., and H.M.; Investigation: N.H., M.H., and H.M.; Resources: M.H.; Data curation: H. Sasaki, N.H.; Writing - original draft: H. Sasaki, N.H.; Writing - review & editing: N.H., M.H., H. Shimojo, H. Sasaki; Visualization: N.H.; Supervision: M.H., H. Shimojo, H. Sasaki; Project administration: H. Sasaki; Funding acquisition: N.H., M.H., H.M., H. Shimojo, H. Sasaki

## Data availability

All relevant data can be found within the article and its supplementary information.

## References

Allegre, N., Chauveau, S., Dennis, C., Renaud, Y., Meistermann, D., Estrella, L.V., Pouchin, P., Cohen-Tannoudji, M., David, L., Chazaud, C., 2022. NANOG initiates epiblast fate through the coordination of pluripotency genes expression. Nature communications 13, 3550.

Avilion, A.A., Nicolis, S.K., Pevny, L.H., Perez, L., Vivian, N., Lovell-Badge, R., 2003. Multipotent cell lineages in early mouse development depend on SOX2 function. Genes & development 17, 126–140.

Behringer, R., Gertsenstein, M., Nagy, K.V., Nagy, A., 2014. Manipulating the mouse embryo : a laboratory manual, Fourth edition. ed. Cold Spring Harbor Laboratory Press, Cold Spring Harbor, New York.

Bessonnard, S., De Mot, L., Gonze, D., Barriol, M., Dennis, C., Goldbeter, A., Dupont, G., Chazaud, C., 2014. Gata6, Nanog and Erk signaling control cell fate in the inner cell mass through a tristable regulatory network. Development (Cambridge, England) 141, 3637–3648.

Briggs, J.A., Weinreb, C., Wagner, D.E., Megason, S., Peshkin, L., Kirschner, M.W., Klein, A.M., 2018. The dynamics of gene expression in vertebrate embryogenesis at single-cell resolution. Science (New York, N.Y 360, eaar5780.

Chazaud, C., Yamanaka, Y., Pawson, T., Rossant, J., 2006. Early lineage segregation between epiblast and primitive endoderm in mouse blastocysts through the Grb2-MAPK pathway. Developmental cell 10, 615–624.

Cong, L., Ran, F.A., Cox, D., Lin, S., Barretto, R., Habib, N., Hsu, P.D., Wu, X., Jiang, W., Marraffini, L.A., Zhang, F., 2013. Multiplex genome engineering using CRISPR/Cas systems. Science (New York, N.Y 339, 819–823.

Farrell, J.A., Wang, Y., Riesenfeld, S.J., Shekhar, K., Regev, A., Schier, A.F., 2018. Single-cell reconstruction of developmental trajectories during zebrafish embryogenesis. Science (New York, N.Y 360, eaar3131.

Frankenberg, S., Gerbe, F., Bessonnard, S., Belville, C., Pouchin, P., Bardot, O., Chazaud, C., 2011. Primitive endoderm differentiates via a three-step mechanism involving Nanog and RTK signaling. Developmental cell 21, 1005–1013.

Frum, T., Watts, J.L., Ralston, A., 2019. TEAD4, YAP1 and WWTR1 prevent the premature onset of pluripotency prior to the 16-cell stage. Development (Cambridge, England) 146.

Guo, G., Huss, M., Tong, G.Q., Wang, C., Li Sun, L., Clarke, N.D., Robson, P., 2010. Resolution of cell fate decisions revealed by single-cell gene expression analysis from zygote to blastocyst. Developmental cell 18, 675–685.

Hashimoto, M., Sasaki, H., 2019. Epiblast Formation by TEAD-YAP-Dependent Expression of Pluripotency Factors and Competitive Elimination of Unspecified Cells. Developmental cell 50, 139–154 e135.

Hashimoto, M., Yamashita, Y., Takemoto, T., 2016. Electroporation of Cas9 protein/sgRNA into early pronuclear zygotes generates non-mosaic mutants in the mouse. Developmental biology 418, 1–9.

Hochedlinger, K., Yamada, Y., Beard, C., Jaenisch, R., 2005. Ectopic expression of Oct-4 blocks progenitor-cell differentiation and causes dysplasia in epithelial tissues. Cell 121, 465–477.

Jacquemin, P., Hwang, J.J., Martial, J.A., Dolle, P., Davidson, I., 1996. A novel family of developmentally regulated mammalian transcription factors containing the TEA/ATTS DNA binding domain. The Journal of biological chemistry 271, 21775–21785.

Jacquemin, P., Sapin, V., Alsat, E., Evain-Brion, D., Dolle, P., Davidson, I., 1998. Differential expression of the TEF family of transcription factors in the murine placenta and during differentiation of primary human trophoblasts in vitro. Dev Dyn 212, 423–436.

Kaneko, K.J., DePamphilis, M.L., 1998. Regulation of gene expression at the beginning of mammalian development and the TEAD family of transcription factors. Dev Genet 22, 43–55.

Kang, M., Piliszek, A., Artus, J., Hadjantonakis, A.K., 2013. FGF4 is required for lineage restriction and salt-and-pepper distribution of primitive endoderm factors but not their initial expression in the mouse. Development (Cambridge, England) 140, 267–279.

Kastan, N., Gnedeva, K., Alisch, T., Petelski, A.A., Huggins, D.J., Chiaravalli, J., Aharanov, A., Shakked, A., Tzahor, E., Nagiel, A., Segil, N., Hudspeth, A.J., 2021. Small-molecule inhibition of Lats kinases may promote Yap-dependent proliferation in postmitotic mammalian tissues. Nature communications 12, 3100.

Kawakami, K., Takeda, H., Kawakami, N., Kobayashi, M., Matsuda, N., Mishina, M., 2004. A transposon-mediated gene trap approach identifies developmentally regulated genes in zebrafish. Developmental cell 7, 133–144.

Koike-Yusa, H., Li, Y., Tan, E.P., Velasco-Herrera, M.D.C., Yusa, K., 2014. Genome-wide recessive genetic screening in mammalian cells with a lentiviral CRISPR-guide RNA library. Nature biotechnology 32, 267–273.

Krawchuk, D., Honma-Yamanaka, N., Anani, S., Yamanaka, Y., 2013. FGF4 is a limiting factor controlling the proportions of primitive endoderm and epiblast in the ICM of the mouse blastocyst. Developmental biology 384, 65–71.

Kurimoto, K., Yabuta, Y., Ohinata, Y., Ono, Y., Uno, K.D., Yamada, R.G., Ueda, H.R., Saitou, M., 2006. An improved single-cell cDNA amplification method for efficient high-density oligonucleotide microarray analysis. Nucleic Acids Res 34, e42.

Kurotaki, Y., Hatta, K., Nakao, K., Nabeshima, Y., Fujimori, T., 2007. Blastocyst axis is specified independently of early cell lineage but aligns with the ZP shape. Science (New York, N.Y 316, 719–723.

Li, L., Lai, F., Hu, X., Liu, B., Lu, X., Lin, Z., Liu, L., Xiang, Y., Frum, T., Halbisen, M.A., Chen, F., Fan, Q., Ralston, A., Xie, W., 2023. Multifaceted SOX2-chromatin interaction underpins pluripotency progression in early embryos. Science (New York, N.Y 382, eadi5516.

Li, M., Belmonte, J.C., 2017. Ground rules of the pluripotency gene regulatory network. Nature reviews 18, 180–191.

Macarthur, B.D., Ma’ayan, A., Lemischka, I.R., 2009. Systems biology of stem cell fate and cellular reprogramming. Nature reviews. Molecular cell biology 10, 672–681.

Mahoney, W.M., Jr., Hong, J.H., Yaffe, M.B., Farrance, I.K., 2005. The transcriptional co-activator TAZ interacts differentially with transcriptional enhancer factor-1 (TEF-1) family members. The Biochemical journal 388, 217–225.

Messerschmidt, D.M., Kemler, R., 2010. Nanog is required for primitive endoderm formation through a non-cell autonomous mechanism. Developmental biology 344, 129–137.

Mitsui, K., Tokuzawa, Y., Itoh, H., Segawa, K., Murakami, M., Takahashi, K., Maruyama, M., Maeda, M., Yamanaka, S., 2003. The homeoprotein Nanog is required for maintenance of pluripotency in mouse epiblast and ES cells. Cell 113, 631–642.

Morris, S.A., Teo, R.T., Li, H., Robson, P., Glover, D.M., Zernicka-Goetz, M., 2010. Origin and formation of the first two distinct cell types of the inner cell mass in the mouse embryo. Proceedings of the National Academy of Sciences of the United States of America 107, 6364–6369.

Ng, H.H., Surani, M.A., 2011. The transcriptional and signalling networks of pluripotency. Nat Cell Biol 13, 490–496.

Nichols, J., Silva, J., Roode, M., Smith, A., 2009. Suppression of Erk signalling promotes ground state pluripotency in the mouse embryo. Development (Cambridge, England) 136, 3215–3222.

Nishioka, N., Inoue, K., Adachi, K., Kiyonari, H., Ota, M., Ralston, A., Yabuta, N., Hirahara, S., Stephenson, R.O., Ogonuki, N., Makita, R., Kurihara, H., Morin-Kensicki, E.M., Nojima, H., Rossant, J., Nakao, K., Niwa, H., Sasaki, H., 2009. The Hippo signaling pathway components Lats and Yap pattern Tead4 activity to distinguish mouse trophectoderm from inner cell mass. Developmental cell 16, 398–410.

Nishioka, N., Yamamoto, S., Kiyonari, H., Sato, H., Sawada, A., Ota, M., Nakao, K., Sasaki, H., 2008. Tead4 is required for specification of trophectoderm in pre-implantation mouse embryos. Mechanisms of development 125, 270–283.

Niwa, H., Toyooka, Y., Shimosato, D., Strumpf, D., Takahashi, K., Yagi, R., Rossant, J., 2005. Interaction between Oct3/4 and Cdx2 determines trophectoderm differentiation. Cell 123, 917–929.

Niwayama, R., Moghe, P., Liu, Y.J., Fabreges, D., Buchholz, F., Piel, M., Hiiragi, T., 2019. A Tug-of-War between Cell Shape and Polarity Controls Division Orientation to Ensure Robust Patterning in the Mouse Blastocyst. Developmental cell 51, 564–574 e566.

Ohnishi, Y., Huber, W., Tsumura, A., Kang, M., Xenopoulos, P., Kurimoto, K., Oles, A.K., Arauzo-Bravo, M.J., Saitou, M., Hadjantonakis, A.K., Hiiragi, T., 2014. Cell-to-cell expression variability followed by signal reinforcement progressively segregates early mouse lineages. Nat Cell Biol 16, 27–37.

Orkin, S.H., Wang, J., Kim, J., Chu, J., Rao, S., Theunissen, T.W., Shen, X., Levasseur, D.N., 2008. The transcriptional network controlling pluripotency in ES cells. Cold Spring Harbor symposia on quantitative biology 73, 195–202.

Ota, M., Sasaki, H., 2008. Mammalian Tead proteins regulate cell proliferation and contact inhibition as a transcriptional mediator of Hippo signaling. Development (Cambridge, England) 135, 4059–4069.

Otsuka, T., Shimojo, H., Sasaki, H., 2023. Daughter cells inherit YAP localization from mother cells in early preimplantation embryos. Dev Growth Differ 65, 360–369.

Plusa, B., Piliszek, A., 2020. Common principles of early mammalian embryo self-organisation. Development (Cambridge, England) 147.

Plusa, B., Piliszek, A., Frankenberg, S., Artus, J., Hadjantonakis, A.K., 2008. Distinct sequential cell behaviours direct primitive endoderm formation in the mouse blastocyst. Development (Cambridge, England) 135, 3081–3091.

Posfai, E., Petropoulos, S., de Barros, F.R.O., Schell, J.P., Jurisica, I., Sandberg, R., Lanner, F., Rossant, J., 2017. Position- and Hippo signaling-dependent plasticity during lineage segregation in the early mouse embryo. Elife 6.

Rossant, J., Lis, W.T., 1979. Potential of isolated mouse inner cell masses to form trophectoderm derivatives in vivo. Developmental biology 70, 255–261.

Saiz, N., Hadjantonakis, A.-K., 2020. Coordination between patterning and morphogenesis ensures robustness during mouse development. Philos Trans R Soc Lond B Biol Sci 375, 20190562.

Saiz, N., Kang, M., Schrode, N., Lou, X., Hadjantonakis, A.K., 2016a. Quantitative Analysis of Protein Expression to Study Lineage Specification in Mouse Preimplantation Embryos. J Vis Exp, 53654.

Saiz, N., Williams, K.M., Seshan, V.E., Hadjantonakis, A.K., 2016b. Asynchronous fate decisions by single cells collectively ensure consistent lineage composition in the mouse blastocyst. Nature communications 7, 13463.

Sato, Y., Kasai, T., Nakagawa, S., Tanabe, K., Watanabe, T., Kawakami, K., Takahashi, Y., 2007. Stable integration and conditional expression of electroporated transgenes in chicken embryos. Developmental biology 305, 616–624.

Schneider, C.A., Rasband, W.S., Eliceiri, K.W., 2012. NIH Image to ImageJ: 25 years of image analysis. Nature methods 9, 671–675.

Schrode, N., Saiz, N., Di Talia, S., Hadjantonakis, A.K., 2014. GATA6 levels modulate primitive endoderm cell fate choice and timing in the mouse blastocyst. Developmental cell 29, 454–467.

Silva, J., Nichols, J., Theunissen, T.W., Guo, G., van Oosten, A.L., Barrandon, O., Wray, J., Yamanaka, S., Chambers, I., Smith, A., 2009. Nanog is the gateway to the pluripotent ground state. Cell 138, 722–737.

Spindle, A.I., 1978. Trophoblast regeneration by inner cell masses isolated from cultured mouse embryos. The Journal of experimental zoology 203, 483–489.

Strumpf, D., Mao, C.A., Yamanaka, Y., Ralston, A., Chawengsaksophak, K., Beck, F., Rossant, J., 2005. Cdx2 is required for correct cell fate specification and differentiation of trophectoderm in the mouse blastocyst. Development (Cambridge, England) 132, 2093–2102.

Sun, X., Ren, Z., Cun, Y., Zhao, C., Huang, X., Zhou, J., Hu, R., Su, X., Ji, L., Li, P., Mak, K.L.K., Gao, F., Yang, Y., Xu, H., Ding, J., Cao, N., Li, S., Zhang, W., Lan, P., Sun, H., Wang, J., Yuan, P., 2020. Hippo-YAP signaling controls lineage differentiation of mouse embryonic stem cells through modulating the formation of super-enhancers. Nucleic Acids Res 48, 7182–7196.

Tamm, C., Kadekar, S., Pijuan-Galito, S., Anneren, C., 2016. Fast and Efficient Transfection of Mouse Embryonic Stem Cells Using Non-Viral Reagents. Stem Cell Rev Rep 12, 584–591.

Vassilev, A., Kaneko, K.J., Shu, H., Zhao, Y., DePamphilis, M.L., 2001. TEAD/TEF transcription factors utilize the activation domain of YAP65, a Src/Yes-associated protein localized in the cytoplasm. Genes & development 15, 1229–1241.

Wagner, D.E., Weinreb, C., Collins, Z.M., Briggs, J.A., Megason, S.G., Klein, A.M., 2018. Single-cell mapping of gene expression landscapes and lineage in the zebrafish embryo. Science (New York, N.Y 360, 981–987.

White, M.D., Zenker, J., Bissiere, S., Plachta, N., 2018. Instructions for Assembling the Early Mammalian Embryo. Developmental cell 45, 667–679.

Wicklow, E., Blij, S., Frum, T., Hirate, Y., Lang, R.A., Sasaki, H., Ralston, A., 2014. HIPPO pathway members restrict SOX2 to the inner cell mass where it promotes ICM fates in the mouse blastocyst. PLoS Genet 10, e1004618.

Xenopoulos, P., Kang, M., Puliafito, A., Di Talia, S., Hadjantonakis, A.K., 2015. Heterogeneities in Nanog Expression Drive Stable Commitment to Pluripotency in the Mouse Blastocyst. Cell reports.

Yagi, R., Kohn, M.J., Karavanova, I., Kaneko, K.J., Vullhorst, D., Depamphilis, M.L., Buonanno, A., 2007. Transcription factor TEAD4 specifies the trophectoderm lineage at the beginning of mammalian development. Development (Cambridge, England) 134, 3827–3836.

Yamanaka, Y., Lanner, F., Rossant, J., 2010. FGF signal-dependent segregation of primitive endoderm and epiblast in the mouse blastocyst. Development (Cambridge, England) 137, 715–724.

Young, R.A., 2011. Control of the embryonic stem cell state. Cell 144, 940–954.

Yusa, K., Zhou, L., Li, M.A., Bradley, A., Craig, N.L., 2011. A hyperactive piggyBac transposase for mammalian applications. Proceedings of the National Academy of Sciences of the United States of America 108, 1531–1536.

Zhang, H.T., Hiiragi, T., 2018. Symmetry Breaking in the Mammalian Embryo. Annual review of cell and developmental biology 34, 405–426.

Zhao, B., Wei, X., Li, W., Udan, R.S., Yang, Q., Kim, J., Xie, J., Ikenoue, T., Yu, J., Li, L., Zheng, P., Ye, K., Chinnaiyan, A., Halder, G., Lai, Z.C., Guan, K.L., 2007. Inactivation of YAP oncoprotein by the Hippo pathway is involved in cell contact inhibition and tissue growth control. Genes & development 21, 2747–2761.

Zhao, B., Ye, X., Yu, J., Li, L., Li, W., Li, S., Yu, J., Lin, J.D., Wang, C.Y., Chinnaiyan, A.M., Lai, Z.C., Guan, K.L., 2008. TEAD mediates YAP-dependent gene induction and growth control. Genes & development 22, 1962–1972.

Zhu, M., Zernicka-Goetz, M., 2020. Principles of Self-Organization of the Mammalian Embryo. Cell 183, 1467–1478.

