## Supplemental Figure for "Fate specification triggers a positive feedback loop of TEAD–YAP and NANOG to promote epiblast formation in preimplantation embryos"

### Supplemental Figure 1

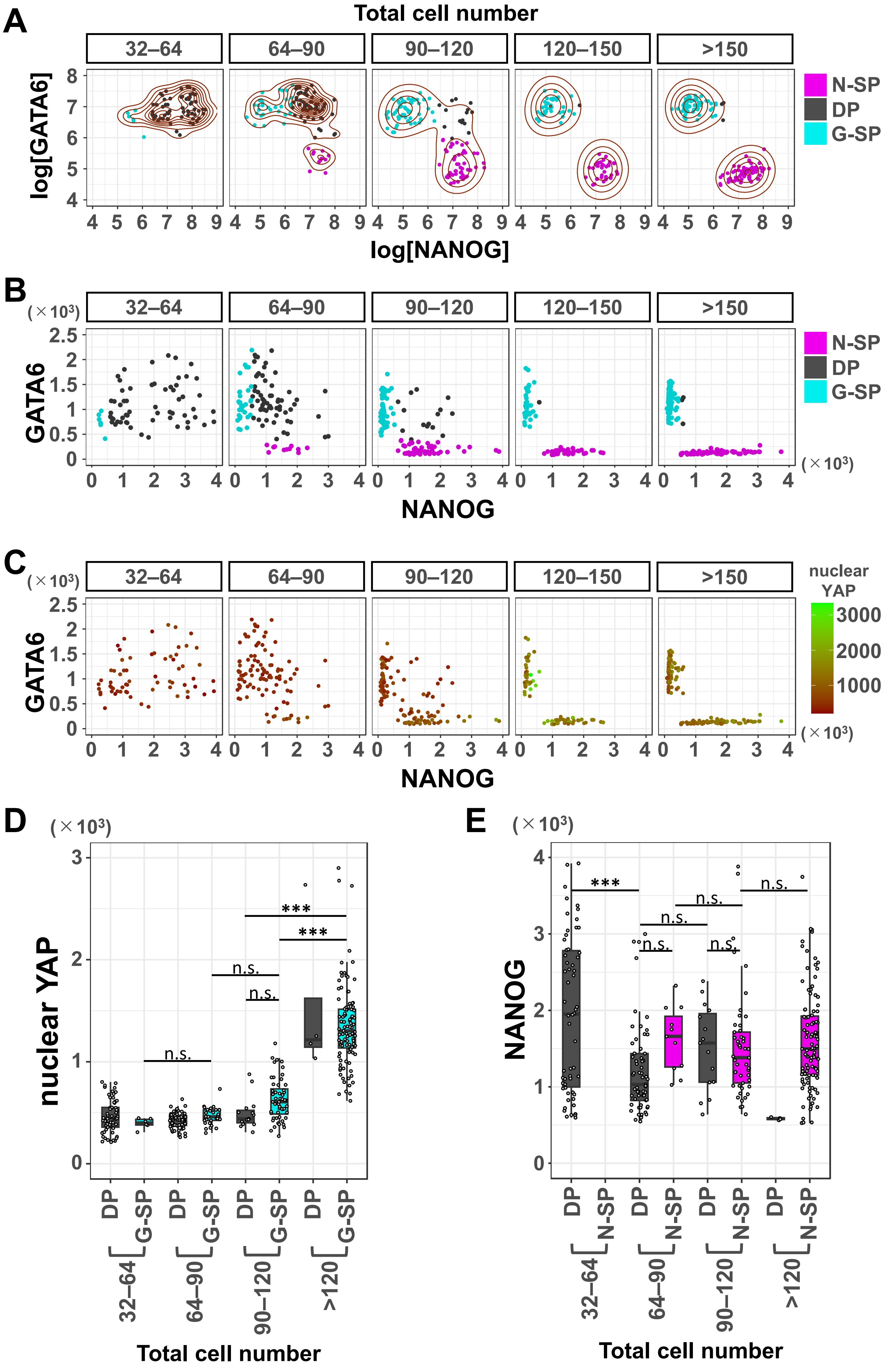

**Supplementary Figure 1.**

**Quantification of NANOG, GATA6, and nuclear YAP signals during blastocyst stages.**

(A) Scatter plots for fluorescence intensity levels of GATA6 and NANOG in individual ICM cells of embryos collected at sequential stages, as in Fig. 1A, represented as logarithm. Cell identity was assigned using a k-means clustering approach. This cell identity was used for all analyses. Contour lines are provided as density estimators. Color coding is indicated. Number of cells analyzed for each stage of the graphs shown in (A) – (C) is as follows: 32–64 cells (n = 67), 64–90 cells (n = 102), 90–120 cells (n = 116), 120–150 cells (n = 73), and >150 cells (n = 138).

(B) Scatter plots for fluorescence intensity levels of GATA6 and NANOG in individual ICM cells of embryos collected at sequential stages, as in Fig. 1A. Values shown in linear scale after inverting the log-transformed, corrected values for each ICM cell. Color coding is indicated.

(C) Scatter plots for fluorescence intensity levels of GATA6 and NANOG with graded colors indicating nuclear YAP signal levels in individual ICM cells of embryos.

(D) Boxplots showing nuclear YAP signal levels in DP and G-SP cells shown in Fig. 1A. Each dot represents the nuclear YAP signal level of a cell. Sample number analyzed for each category is as follows: 32–64 cells, DP (n = 61), G-SP (n = 6); 64–90 cells, DP (n = 61), G-SP (n = 30); 90–120 cells, DP (n = 15), G-SP (n = 52), >120 cells, DP (n = 4), G-SP (n = 104). *p*-values were determined through two-way ANOVA followed by Tukey's post hoc multiple comparison test. $^{***}p < 0.001$, n.s. not significant.

(E) Boxplots showing NANOG signal levels in DP and N-SP cells shown in Fig. 1A. Each dot represents the NANOG signal level of a cell. Sample numbers and statistical analysis are the same as in Fig. 1C.

### Supplemental Figure 2

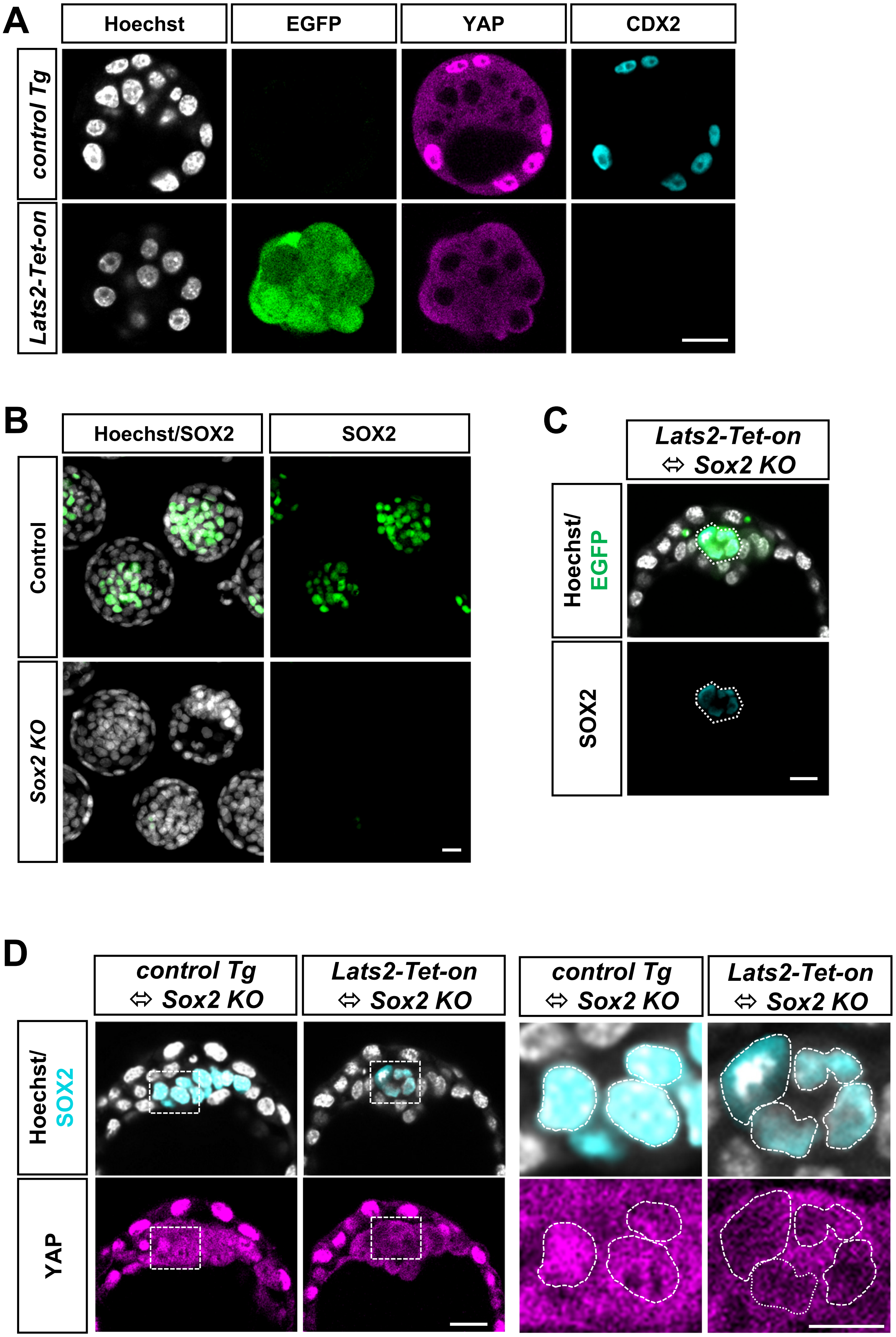

**Supplementary Figure 2.**

**Characterization of *Lats2-Tet-on* and *Sox2* KO embryos.**

(A) Representative immunofluorescence images of EGFP, YAP and CDX2 expression in *Lats2-Tet-on* and *Control Tg* embryos. Scale bar represents 20 µm.

(B) Representative immunofluorescence images of SOX2 expression in *Sox2 KO* embryos. Z-stack images are shown. Scale bar represents 20 µm.

(C) Representative immunofluorescence images of EGFP, and SOX2 expression in *Lats2-Tet-on* ⇔ *Sox2 KO* embryos. Scale bar represents 20 µm. Hoechst and EGFP signals are shown in grayscale and green, respectively. Dashed line indicates EPI. Scale bar represents 20 µm.

(D) Enlarged view of Figure 2C. The left four panels show the same sections as in Figure 2C. SOX2 and Hoechst signals are merged in the upper panels. The boxed areas are enlarged in the right four panels. The dashed lines indicate nuclei. The scale bars represent 20 µm in the left panels and 10 µm in the right panels.

### Supplemental Figure 3

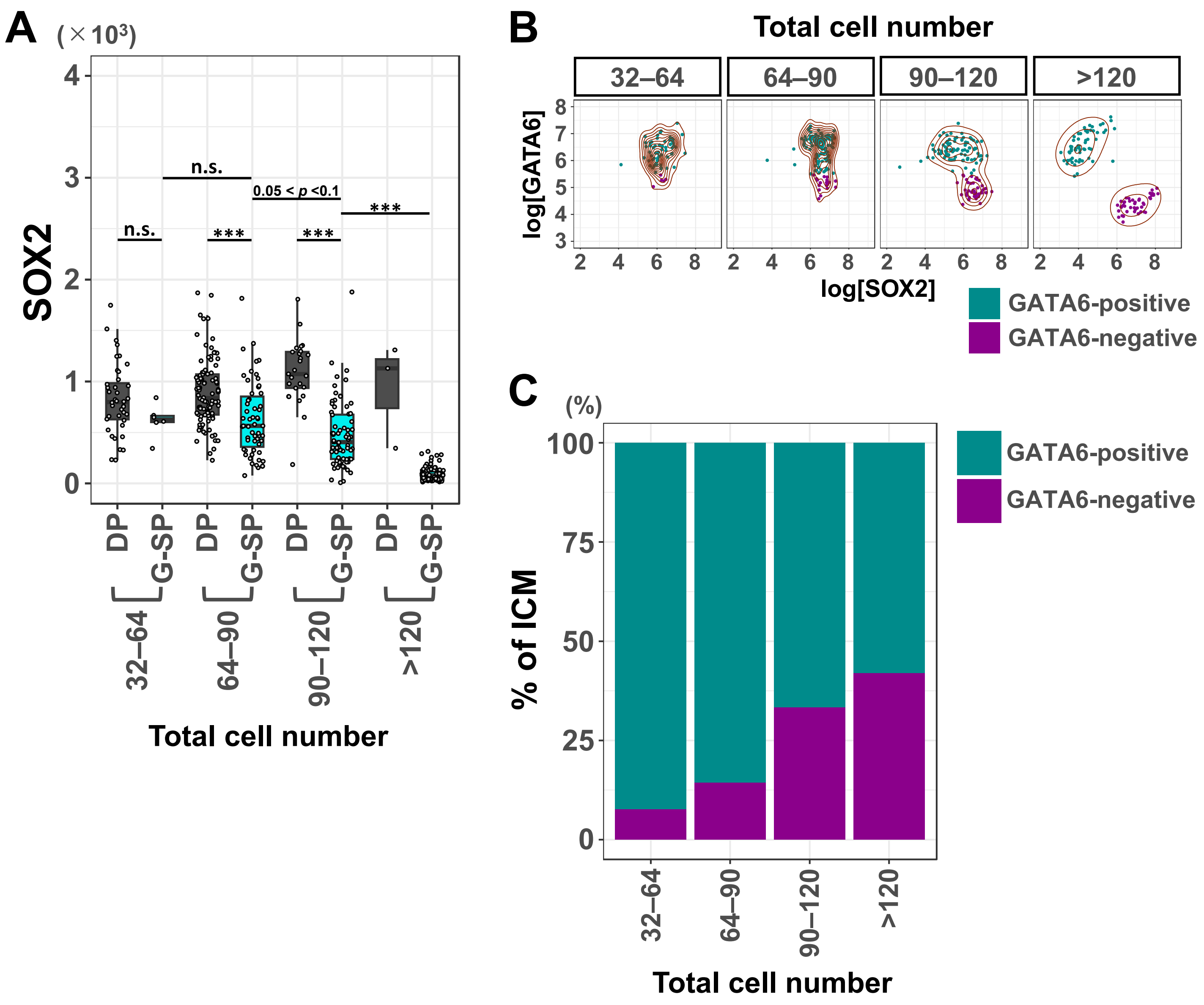


**Supplementary Figure 3.**
**Quantification of GATA6 and SOX2 signals.**
(A) Boxplots showing SOX2 signal levels in DP and G-SP cells shown in Fig. 4A. Each dot represents the SOX2 signal level of a cell. Sample number analyzed for each category is as follows: 32–64 cells, DP (n = 40), G-SP (n = 6); 64–90 cells, DP (n = 92), G-SP (n = 59); 90–120 cells, DP (n = 25), G-SP (n = 72), >120 cells, DP (n = 3), G-SP (n = 81). *p*-values were determined through two-way ANOVA followed by Tukey's post hoc multiple comparison test. ***$p < 0.001$.
(B) Scatter plots for fluorescence intensity levels of GATA6 and SOX2 in individual ICM cells of embryos collected at sequential stages, as in Fig. 4C, represented as logarithm. Cell identity was assigned using a k-means clustering approach. Contour lines are provided as density estimators. Number of cells analyzed for each stage is as follows: 32–64 cells (n = 676), 64–90 cells (n = 133), 90–120 cells (n = 129), and >120 (n = 105). Color coding is indicated.
(C) Average ICM composition, as in Fig. 4C, binned in the developmental stages indicated. Color coding is indicated. Number of cells analyzed for each stage is as follows: 32–64 cells (n = 66), 64–90 cells (n = 133), 90–120 cells (n = 129) and >120 cells (n = 105).

### Supplemental Figure 4

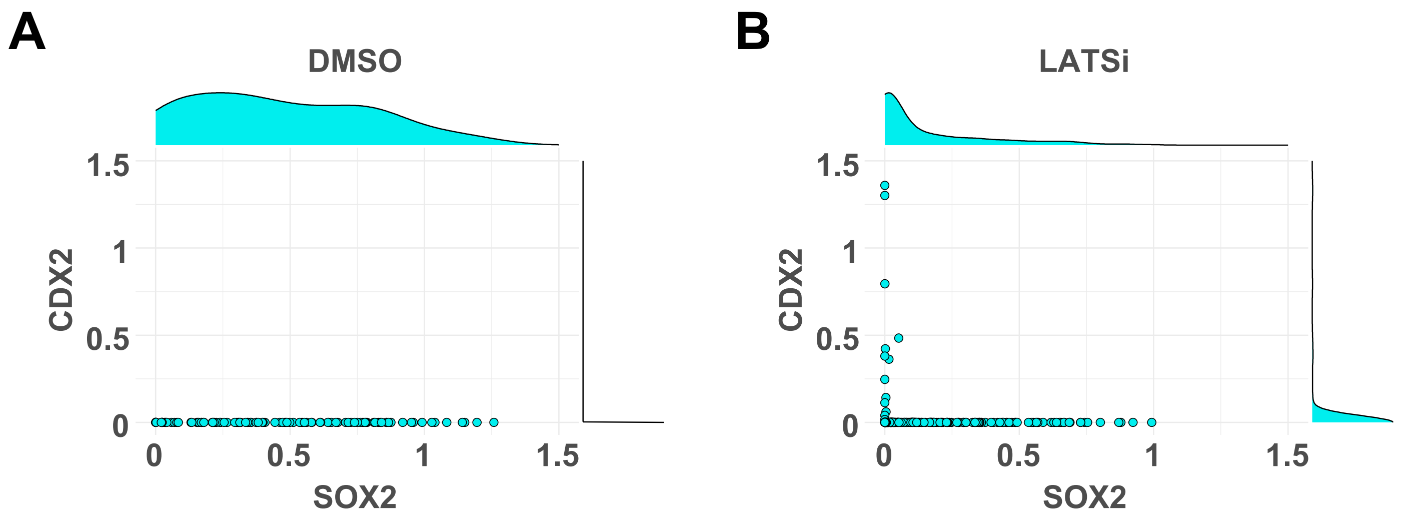


**Supplementary Figure 4.**
**Distribution of CDX2 and SOX2 signals in embryos treated with LATSi for 12 hours from early mid-blastocyst stage.**
(A) Dot plot and half-violin plots showing distributions of CDX2 and SOX2 signal levels in the ICM of the embryos treated with DMSO (n = 103) shown in Fig. 5F–H.
(B) Dot plot and half-violin plots showing distributions of CDX2 and SOX2 signal levels in the ICM of the embryos treated with LATSi (n = 277) shown in Fig. 5F–H.

### Supplemental Figure 5

**A**

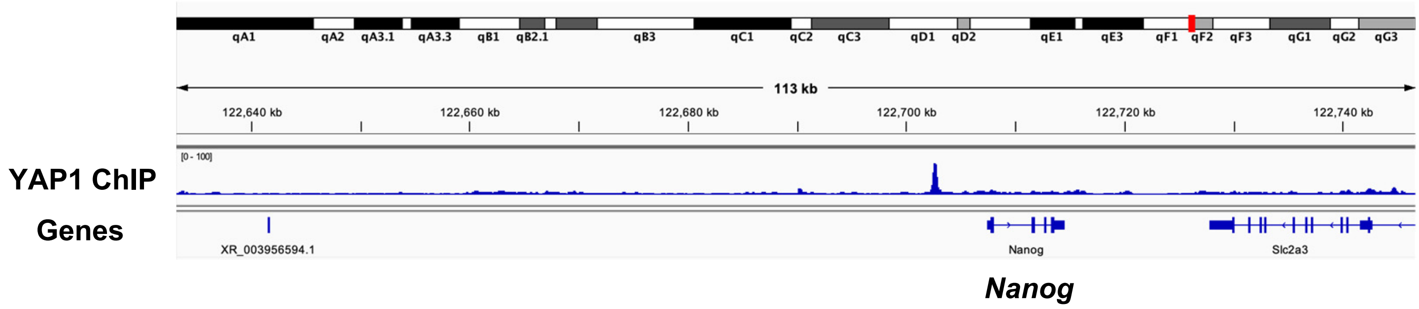


**B**

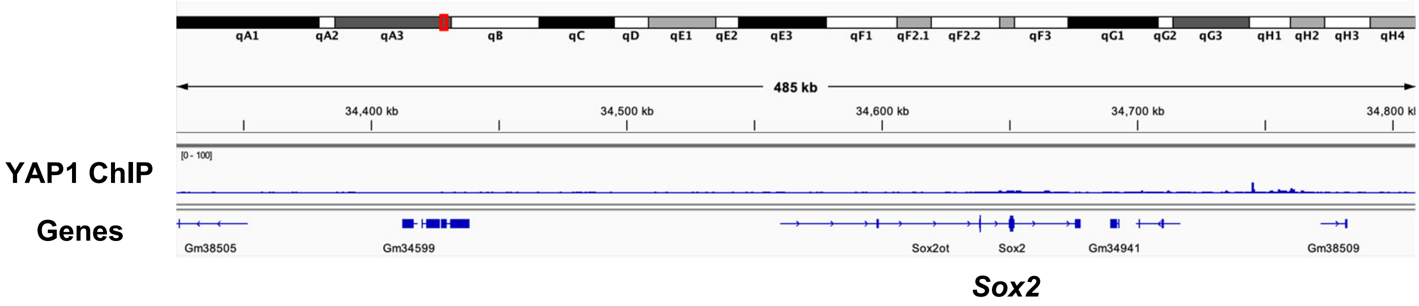


**Supplementary Figure 5.**
**YAP1 binding sites around *Nanog* and *Sox2* in ES cells.**
Distribution of YAP1 ChIP-seq peaks in ES cells around *Nanog* (A) and *Sox2* (B). ChIP-seq data of GSM4291129 (Sun et al., 2020) was visualized with Integrative Genomics Viewer.
